# Thermosensitive alternative splicing senses and mediates temperature adaptation in *Drosophila*

**DOI:** 10.1101/503409

**Authors:** Naveh Evantal, Ane Martin Anduaga, Osnat Bartok, Ines Lucía Patop, Ron Weiss, Sebastian Kadener

## Abstract

Circadian rhythms are generated by the cyclic transcription, translation and degradation of clock genes, including *timeless* (*tim*). Currently, little is known about the mechanisms by which the circadian clock senses and adapts to temperature changes. Here we show that temperature dramatically changes the splicing pattern of *tim*. We found that at 18°C TIM protein levels are diminished due to the induction of two cold-specific splicing isoforms (*tim-cold* and *tim-short&cold*). At 29°C, another isoform, *tim-Medium* is strongly upregulated. We found that this isoform switching mechanism allows flies to regulate the levels and activity of TIM by setting miRNA-dependent thresholds for expression as well as by expressing isoforms with specific functions. Flies in which the production of *tim-short&cold* is abrogated display altered patterns of locomotor activity and altered *tim* expression. Interestingly, the introns of *tim* carry the information for the temperature sensitivity, suggesting that *tim* splicing *per se* is the temperature sensor.

## INTRODUCTION

Circadian rhythms organize most physiological and behavioral processes to 24 hours cycles (Allada and Chung, 2010; Pilorz et al., 2018). The current model postulates that circadian clocks keep time through a complex transcriptional-translational negative feedback loop that takes place in the so-called “clock cells” (Helfrich-Forster, 2003; Shafer et al., 2006). Each one of these clock cells has been proposed to function autonomously (each cell is its own oscillator) (Dissel et al., 2014; Yoshii et al., 2012). In *Drosophila*, the master regulators CLOCK (CLK) and CYCLE (CYC) activate the circadian system by promoting rhythmic transcription of several key clock genes. Products of three of these genes, PERIOD (PER), TIMELESS (TIM), and CLOCKWORK ORANGE (CWO) repress CLK-CYC mediated transcription in an oscillatory manner (Hardin and Panda, 2013; Zhou J, 2016). These cycles of transcriptional activation and repression lead to 24-hour molecular oscillations, which ultimately generate behavioral rhythms. In addition to transcriptional control, post-transcriptional and post-translational regulatory processes play essential roles in circadian timekeeping (Hardin and Panda, 2013; Harms et al., 2004; Kojima et al., 2011; Lim and Allada, 2013). The transcriptional repressors PER and TIM are post-translationally modified, and the modification status as well as the rates with which these modifications take place have a significant influence on their degradation rates (Hardin and Panda, 2013; Ozkaya and Rosato, 2012). Modification of PER may additionally influence its transcriptional repressor activity or the timing of its activity (Hardin and Panda, 2013; Ozkaya and Rosato, 2012).

Circadian clocks are extraordinarily robust systems; they are able to keep time accurately without timing cues. In addition, and despite their biochemical nature, they are resilient to large variations in environmental conditions. The robustness of the circadian system is likely the result of multiple layers of regulation that assure accurate timekeeping and buffering of stochastic changes into the molecular clockwork. These levels of regulation are physically and/or functionally interconnected and, importantly, extend even beyond the single-cell level. Circadian neurons in the brain are organized in a network that is believed to synchronize the individual neuronal oscillators thereby contributing to a coherent and robust behavioral output (Allada and Chung, 2010; Mezan et al., 2016; Peng et al., 2003; Schlichting et al., 2016; Stoleru et al., 2004; Weiss et al., 2014). The PDF neuropeptide, main neuromodulator of the circadian neuronal network, is expressed in a small subset of these neurons. PDF is essential for normal circadian activity patterns in light:dark (LD) cycles and for persistent circadian rhythms in constant darkness (DD) (Im et al., 2011; Peng et al., 2003; Taghert and Shafer, 2006). Circadian clocks are also remarkably plastic systems. For example, organisms can quickly adjust to different light and temperature regimes (Roessingh et al., 2015; Wolfgang et al., 2013). The plasticity of the circadian clock results from the existence of very efficient input pathways that can convey the external signals into the core oscillator machinery (Bartok et al., 2013; Ogueta et al., 2018).

*Timeless* is at the crossroad of both the robustness and plasticity of the circadian clock. *tim* is a core circadian oscillator component and mutations in *tim* coding sequence lead to flies with short, long, or no circadian rhythms (Myers et al., 1995; Rothenfluh et al., 2000; Sehgal et al., 1994). TIM stabilizes PER, which is also essential for circadian rhythmicity, and this complex represses CLK-mediated transcription in an oscillatory manner (Hall, 2003). Importantly, the stoichiometric relationship between PER, CLK and TIM is tightly controlled and probably the major regulator of circadian period (Fathallah-Shaykh et al., 2009; Hardin, 2011; Kadener et al., 2008; Yu et al., 2006). In addition, TIM is the key factor for communicating external information to the central core oscillator. For example, upon light stimulus, the protein encoded by *cryptochrome*, CRY, binds to TIM and promotes TIM degradation through the ubiquitin ligase *jetlag*, which results in phase advances or delays of the circadian oscillator (Emery et al., 1998; Koh et al., 2006; Lin et al., 2001). Last but not least, PDF signalling has been proposed to synchronize the circadian clock by regulating TIM degradation (Li et al., 2014; Seluzicki et al., 2014), suggesting that TIM is the key pathway for conveying external information to the circadian system at the molecular level.

Temperature has diverse effects on the *Drosophila* circadian system (Afik et al., 2017; Kidd et al., 2015). Flies show temperature-dependent changes in the distribution of daily locomotor activity, named seasonal adaptation (Low et al., 2008; Majercak et al., 1999). They are also entrained/synchronized by daily temperature cycles (Roessingh et al., 2015), are phase-shifted by temperature pulses or steps (Glaser and Stanewsky, 2005), and can keep 24-hour periods in a wide range of temperatures (temperature compensation (Kidd et al., 2015)). Adaptation of *Drosophila* to different environmental conditions relies on the presence of at least two neuronal circuits: one that controls the morning peak of locomotor activity (M) and another that controls the evening (E) burst of activity (Grima et al., 2004; Stoleru et al., 2004; Stoleru et al., 2005). The environment fine-tunes the activity pattern by altering the timing of these oscillators. In the laboratory, mimicking summer by subjecting the flies to hot temperatures and long photoperiods shift their M activity to pre-dawn and their E activity into the early night (Majercak et al., 1999). In contrast, under shorter day lengths and cooler temperatures, mimicking autumn, the M and E activity components fall closer together and occur around the middle of the day, enabling maximal activity at the warmest temperatures.

Work from the Edery lab in 1999 first linked the splicing of *per* 3’ untranslated region (UTR) to seasonal adaptation (Majercak et al., 1999). The evening advance in locomotor activity at lower temperatures correlates with an advance in the phase of oscillation of *tim* and *per* mRNAs. The authors postulated that this shift of the clock is driven by the splicing of an alternative exon located in *per* 3’ UTR. Further studies revealed that the efficiency of *per* mRNA splicing is regulated not only by temperature but also by the photoperiod (Collins et al., 2004). The effect of light on *per* splicing does not require a functional clock and phospholipase C likely plays a physiological, non-photic role in downregulating the production of the spliced transcripts (Collins et al., 2004). Moreover, in 2008, a new work examined the splicing pattern of *per* in the tropical species *Drosophila yakuba*, which faces no significant seasonal variation in day length or temperature (Ko et al., 2003; Low et al., 2008; Russo et al., 1995). Although the *per* gene of *D. yakuba* has a 3’-terminal intron, this intron is removed over a wide range of temperatures and photoperiods, consistent with the marginal effect of temperature on this tropical species. Despite these findings, it is still not clear how the regulation of the 3’ UTR splicing impacts PER levels or if other transcriptional and/or post-transcriptional events are also responsible for the regulation of activity in response to temperature changes.

Post-transcriptional control by miRNAs and RNA binding proteins has been shown to be important for circadian timekeeping in flies and other organisms (Chen and Rosbash, 2016; Kadener et al., 2009; Lerner et al., 2015; Lim and Allada, 2013; Xue and Zhang, 2018). On the other hand, only more recently a few reports show a role of alternative splicing on circadian timekeeping (Bartok et al., 2013; Petrillo et al., 2011; Sanchez et al., 2010; Wang et al., 2018). This is surprising, given the prevalence of alternative splicing in the brain and the importance of this process in regulating the amount and type of mRNAs generated from a given locus (Baralle and Giudice, 2017; Sanchez et al., 2011). Alternative mRNA processing is co-transcriptional and usually happens due to the presence of suboptimal processing signals (i.e. splice or cleavage and polyadenylation sites). Many factors can influence alternative splicing including changes in RNA structure, the presence of RNA binding proteins and the elongation rate of RNA Polymerase II (Caceres and Kornblihtt, 2002; de la Mata et al., 2003).

Here we show that temperature dramatically and specifically changes the splicing pattern of *tim*. We found that the lower levels of the canonical TIM protein at 18°C are due to the induction of two cold-specific splicing isoforms (*tim-cold* and *tim-short&cold*). *Tim-cold* encodes a canonical TIM protein but it is under strong miRNA-mediated control as showed by AGO1 IP and *in vivo* luciferase reporters. Moreover, *tim-short&cold* (*tim-sc*) encodes a short TIM isoform which can advance the phase of the circadian clock when overexpressed. Interestingly, most of the changes in *tim* splicing patterns are conserved across *Drosophila* species and correlate well with the capacity of the species to adapt their activity to temperature changes. We follow by using CRISPR to generate flies in which the production of *tim-sc* is abrogated. These flies display altered patterns of locomotor activity at 18 and 25°C as well as altered expression of the remaining *tim* isoforms, demonstrating the importance of *tim-sc* production. Moreover, we showed that the temperature-dependent changes in *tim* alternative splicing are independent of the circadian clock. Furthermore, we could reproduce them in *Drosophila* S2 cells and utilizing splicing minigenes. The latter results strongly suggest that the intronic sequences of *tim* themselves are the temperature sensor for these changes in splicing.

## RESULTS

### Temperature remodels the circadian transcriptome

To determine the effect temperature has on general and circadian gene expression we evaluated gene expression genome-wide of heads of flies at 18°C, 25°C or 29°C. Briefly, we performed 3’ sequencing at 6 different timepoints in flies entrained to 12:12 Light:Dark (LD) cycles at these three temperatures. We identified hundreds of genes displaying oscillating expression over the day (Figure 1A and Table S1). Interestingly, most of the cycling RNAs were temperature specific (Figure 1B, Table S1), with only a few RNAs consistently displaying mRNA oscillations at all the assayed temperatures. The set of oscillating mRNAs was enriched for different Gene Ontology (GO) terms, demonstrating that the circadian program is temperature dependent (Table S2). Interestingly, RNAs oscillating at 18°C display earlier phases than those cycling at 25 or 29°C (Figure 1A and Table S1). This phase advance was even more noticeable on the RNAs oscillating at more than one temperature including the core circadian components (Figure 1C), suggesting that the circadian clock gene expression program is advanced at 18°C. These phase advance mirrors the advance of the evening peak of activity observed at 18°C. Interestingly and despite the clear behavioral differences, we observed smaller phase changes in the flies entrained at 29°C compared to those at 25°C (Figure 1A, S1 and Table S1). Interestingly, our data revealed no major changes in the overall levels of most circadian components when entrained at different temperatures (Figure 1D). As stated above (and previously described (Majercak et al., 1999)), *tim* and *per* mRNA profiles were advanced at 18°C. This phase advance is transcriptional, as can be observed by assessing *tim* and *per* RNAs from nascent (chromatin-bound) RNA (Figure S2). This phase advance in the mRNA level provokes a phase advance notable in TIM and PER protein levels (Majercak et al., 1999).

**Figure 1.**
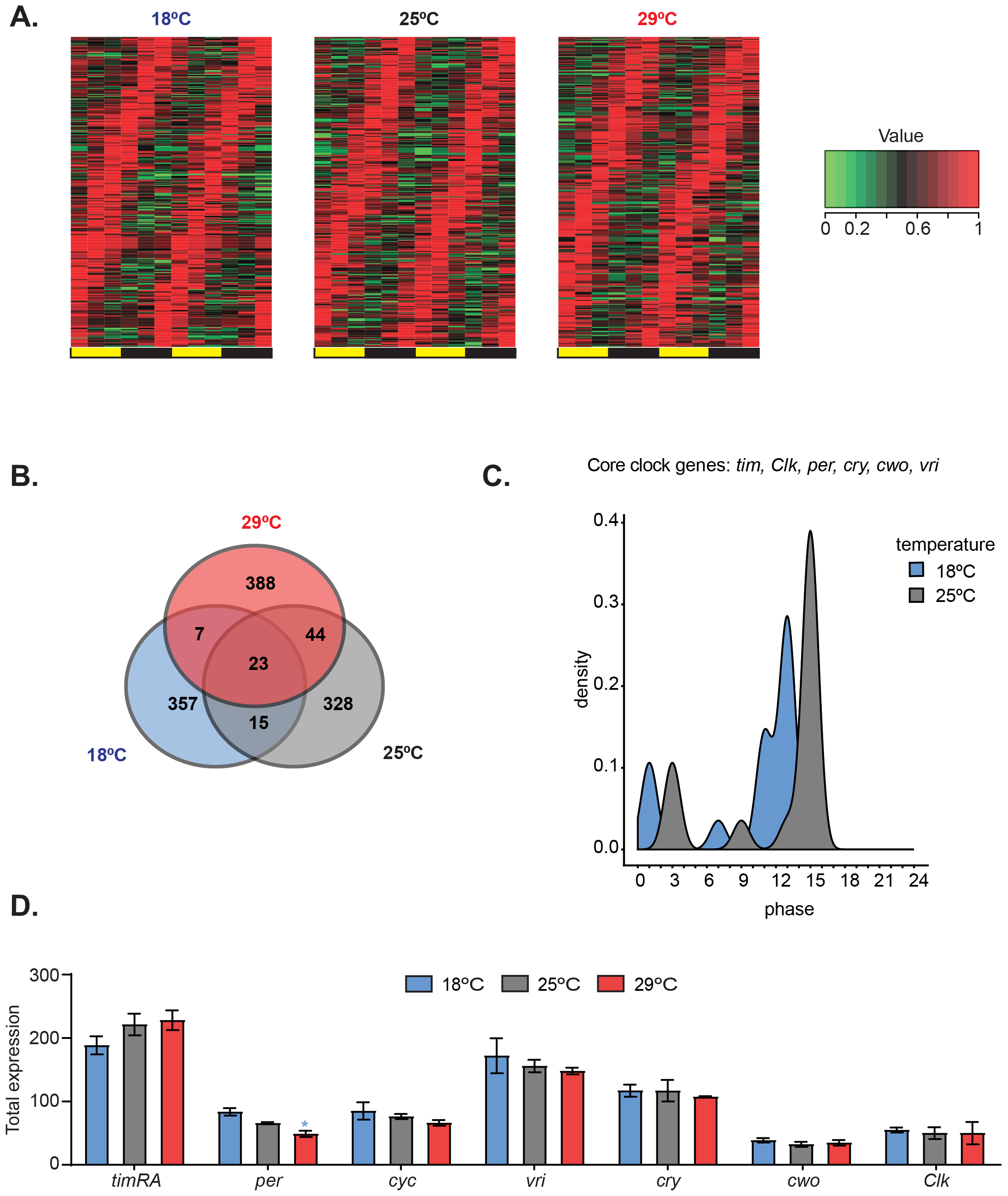
Temperature remodels the circadian transcriptome. **A.** Heat maps of normalized gene expression for genes that display 24hs cycling expression at different temperatures. Flies were entrained at the indicated temperatures for 3 days in 12:12 Light: Dark (LD) conditions. Fly heads were collected every 4 hours in 12 independent timepoints. Cycling expression was assessed as indicated in material and methods. **B.** Venn diagrams indicating the number of oscillating genes at different temperatures. **C.** Density plot showing the phases of core clock genes cycling at both 18°C and 25°C. The blue and grey shades indicate the phase of these genes at 18°C and 25°C respectively. **D.** Total expression of the indicated clock genes in the microarray for each temperature. Only *per* shows a significant difference between 18°C and 29°C.

To complement these data, we analyzed gene expression at the three temperatures utilizing oligonucleotide microarrays from samples that were collected at 6 timepoints but pooled before the gene expression assessment. Pooling the timepoints allows a more precise comparison of average gene expression, avoiding big variations due to timepoints. When examining the Microarray data we were able to identify hundreds of mRNA transcripts that change their expression in response to adaptation to different temperatures. We found changes in both directions, i.e higher or lower at high/low temperatures (Figure S3A). We found a number of Gene Ontology (GO) categories enriched among the differentially expressed genes, spanning a wide range of biological functions (Figure S3B). These results are consistent with previously reported data (Boothroyd et al., 2007a). For example, we observed an increase in expression of ribosomal proteins, genes involved in protein folding and metabolic processes at 29°C. At 18°C we saw increased expression of proteins involved in cuticle development, which is consistent with previous reports showing that cuticle deposition is temperature dependent (Boothroyd et al., 2007a; Ito et al., 2011). These results demonstrate the dramatic effect that temperature has on mRNA steady-state levels and on various cellular and metabolic processes. Overall, the data above demonstrates the eminent role of temperature in global gene expression, and specifically on the circadian molecular and transcriptional network.

### Temperature modulates tim alternative splicing

3’-seq datasets are a powerful tool for examining changes in gene expression. However, they lack the ability to capture many processing events such as alternative splicing and alternative promoter usage. Therefore, we generated whole-transcript polyA^+^ RNA-seq datasets from heads of flies entrained at 18°C, 25°C or 29°C. We found that temperature changes greatly impact the pattern of alternative splicing of *tim* mRNA but not any other clock gene (Figure 2A and data not shown). We identified four major isoforms generated from the *tim* locus (Figure 2B). Except for the canonical transcript, which is generated from an already characterized transcript (referred to from now on as *tim-L*), all other three isoforms changed their expression in a temperature dependent manner. Two of those were more abundant at 18°C compared to 25°C. The first one was previously described as being induced by cold temperatures (*tim-cold*) (Boothroyd et al., 2007a; Wijnen et al., 2006). It includes the last intron of *tim* and has a stop codon at the beginning of the intron, generating a slightly smaller protein than the canonical *tim-L* isoform, but with a longer 3’ UTR. The second cold-enriched isoform, although annotated, was not previously described, and we named it *tim-short&cold* (abbreviated *tim-sc*), as putatively encodes a much smaller protein. In contrast to the other isoforms which are generated by intron retention, *tim-sc* mRNA is generated by the usage of an alternative cleavage and polyadenylation site located with the intron 10 of *tim* which can only be used before this intron is spliced out. The third isoform we identified potentially encodes for a medium sized protein (smaller than those encoded by *tim-L* and *tim-cold* and larger than *tim-sc*) that we named *tim-medium* (abbreviated *tim-M*). This isoform was recently described and named *tim-tiny* (Shakhmantsir et al., 2018). However, to avoid confusion with *tim-sc* (which encodes an even smaller isoform), we will call this isoform *tim-M*. This isoform, is generated by the inclusion of an intron containing a stop codon and shares all downstream exons with *tim-L*, hence presenting the longest 3’ UTR of all isoforms. We found that this isoform is highly expressed in flies entrained to 25°C or 29°C compared to flies kept at 18°C.

**Figure 2.**
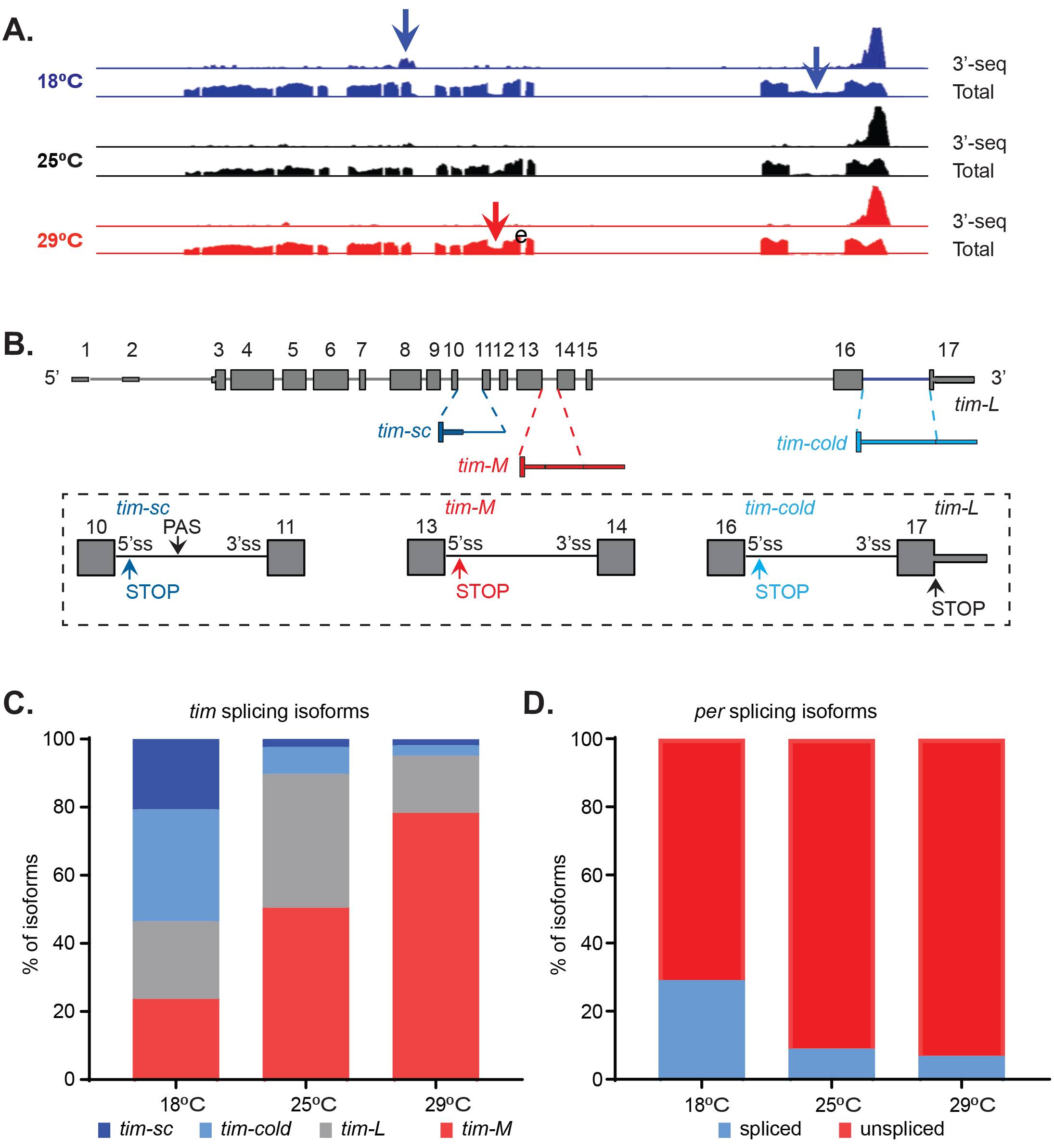
Temperature modulates *tim* alternative splicing. **A.** IGV snapshot of the *timeless* locus region indicating the expression of this gene at 18°C (blue), 25°C (black) and 29°C (red). The presented data includes the aggregated data from the 3’-seq datasets presented in Figure 1 (upper traces) as well as full transcript polyA^+^ RNA seq datasets (lower traces). The latter includes 2 timepoints at 25°C and three timepoints at 18°C and 29°C (see text for details). The arrows indicate the alternative splicing events that are regulated by temperature. **B.** A scheme of the alternatively spliced *tim* isoforms. In grey are constitutive exons, in red sequences found mainly at high temperatures and in blue, sequences found mainly at low temperatures. A zoom on the exons surrounding each non-canonical isoform is represented in the rectangle. Alternative stop codons (STOP) and cleaveage and polyadenylation sites (PAS) in these isoforms are also indicated. **C.** Quantification of the relative amount of *tim-sc* (dark blue), *tim-cold* (light blue), *tim-M* (red) and *tim-L* (grey) isoforms at the three assayed temperatures. To quantify the different isoforms we counted the relative number of spliced junctions specific for each isoform. **D.** Quantification of the relative amount of *per-spliced* (blue) and *per-unspliced* (red) RNAs at the three assayed temperatures performed as in C.

Since three of these isoforms (*tim-L, tim-cold and tim-M*) share the same 3’ end, they are undistinguishable in the 3’-seq datasets. In contrast, we could easily detect a significant increase in the levels of *tim-sc* mRNA at 18°C in this dataset (Figure S4). We also could observe the effect in the whole-transcript datasets, although the differences were more difficult to appreciate due to the underrepresentation of 3’ ends in whole-transcript sequencing. We confirmed the changes detailed above by qRT-PCR using isoform specific primers (Figure S5A). Importantly, the levels of *tim-L* cannot be unequivocally assessed by qPCR as its sequence overlaps with *tim-M*. Temperature-dependent changes in the splicing patterns of *tim* are, at least, as impressive as the previously described changes in the splicing of a small intron in the 3’ UTR of *per* (Majercak et al., 1999).

We next utilized the whole RNA-seq datasets to quantify the relative amount of each of the four isoforms (Figure 2C). We found the change in the distribution of the four isoforms between the different temperatures to be clearly dramatic, from a fairly even distribution at 18°C to no expression of *tim-sc* and *tim-cold* and significant elevation of *tim-M* at 29°C. Surprisingly, we found that *tim-M* constitutes about fifty percent of *tim* mRNAs at 25°C, having a similar, if not higher, expression than the canonical isoform *tim-L*. This suggests the existence of previously unknown aspects of *tim* regulation even in canonical conditions. In comparison to the big changes driven by temperature observed in *tim* alternative splicing, the well-characterized changes in *per* alternative splicing are very modest (Low et al., 2008) (Figure2D). To assess if the observed changes in *tim* RNA processing upon temperature changes also happen in the fly brain, we dissected the brains of flies entrained at 18, 25 and 29°C and measured the levels of each *tim* isoform by qPCR. As observed when utilizing fly heads, entrainment of the flies to lower temperatures (18°C) resulted in a strong increase of *tim-s* and *tim-cold* levels while higher temperatures (29°C) promoted *tim-M* expression (Figure S5B).

### Temperature-specific alternative splicing is conserved across Drosophilae and correlates with temperature adaptation

If changes in the splicing pattern of *tim* are necessary for temperature adaptation, it should be conserved among other *Drosophila* species that adapt to temperature changes. Therefore, we determined both the behavioral and splicing patterns of three additional species, belonging to the *Drosophila* genus at 18, 25 and 29°C. Two of those species (*D.simulans* and *D.Yakuba*) belong, like *D.melanogaster*, to the *Sophophora* subgenus, while the third (*D.virilis*) belongs to the *Drosophila* subgenus. Furthermore, even if they share the same tropic origin, *D.melanogaster* and *D.simulans* have become cosmopolitan, while the distribution of *D.Yakuba* is restricted to Africa (Kuntz and Eisen, 2014; Markow and O’Grady, 2007). *D.virilis*, a Holarctic species, is also geographically temperate. Although all species displayed some degree of behavioral adaptation to temperature, we could detect differences between species when analyzing the ratio between light and dark activity in all three temperatures (Figure 3A). As expected, *D.melanogaster* displays a temperature-dependent gradient in the day/night activity ratio, with highest ratios at 18°C (Figure 3A, left). *D.simulans* showed the same trend of significant decrease in light/dark activity ratios in elevating temperatures, suggesting behavioral adaptation to high and low temperatures (Figure 3A, middle-left; Figure S6A upper left panel and data not shown). In contrast, while adjusting to high temperatures, *D.Yakuba* did not show any significant difference between the ratios of 25°C and 18°C, suggesting reduced adaptational capacity to lower temperatures (Figure 3A, middle-right; Figure S6A middle left panel and data not shown). It should be mentioned that while the ratio at 29°C in *D.virilis* is the same as in the other species, its activity was affected in the opposite trend than the others, decreasing during the dark (Figure 3A, right; Figure S6A lower left panel and data not shown).

**Figure 3.**
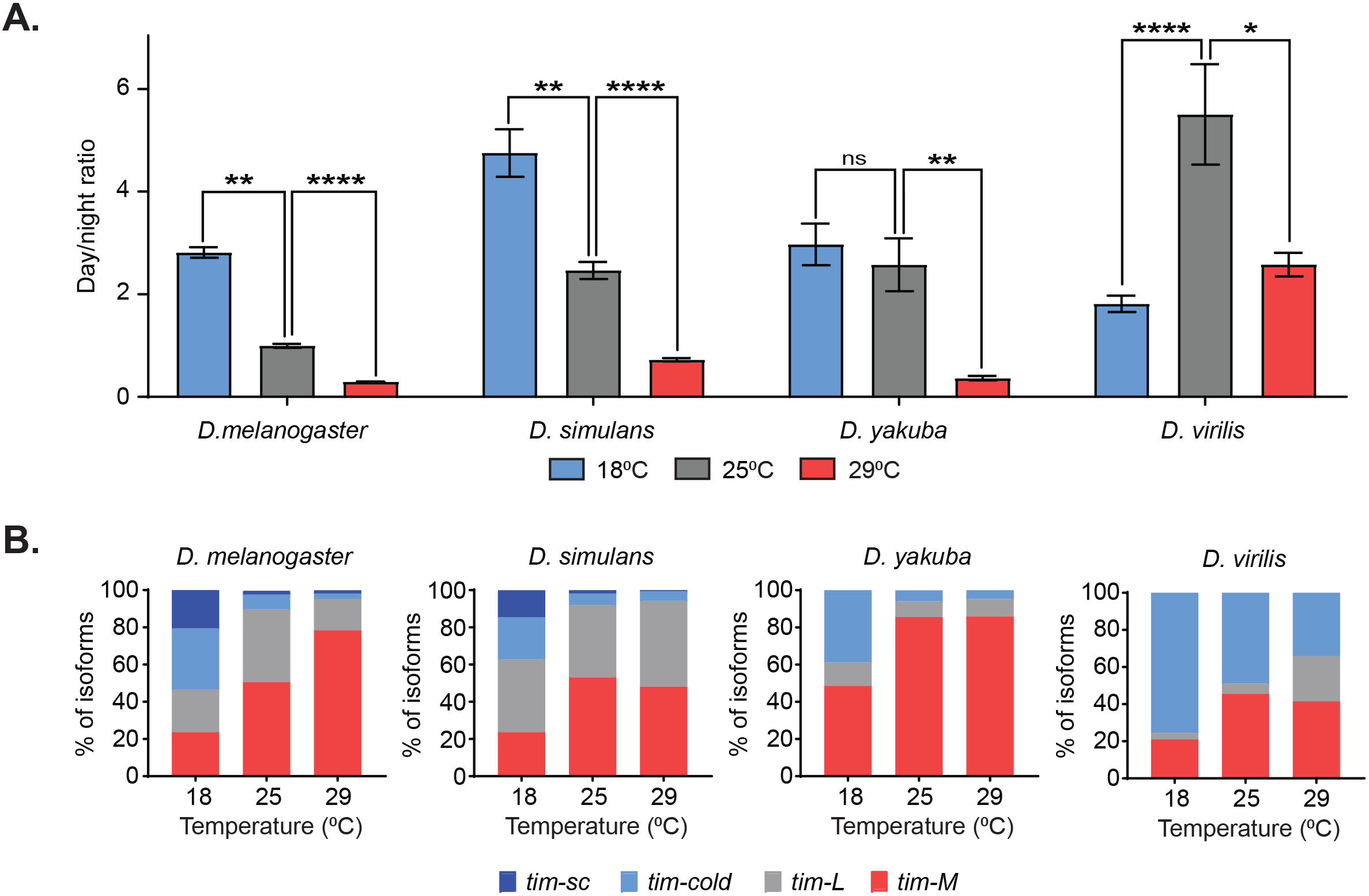
Temperature-specific alternative splicing is conserved across different *Drosophila* species. **A.** Quantification of the ratio between day and night activity counts of locomotor activity for *D. melanogaster, D. simulans, D. yakuba* and *D. virilis* at 18°C (blue), 25°C (grey) or 29°C (red). Each bar represents the ratio between the mean locomotor activities (light and dark) of all four days (N~32). Significance was calculated performing multiple t-test analysis using the Holm-Sidak method.(*, pval<0.05; **, pval<0.01 ***, pval<0.001; ****, pval<0.0001). **B.** Quantification of the relative amount of *tim-sc* (dark blue), *tim-cold* (light blue), *tim-M* (red) and *tim-L* (grey) from whole transcriptome (polyA^+^) mRNA-seq in the species at the indicated temperatures. The ratios between the isoforms were calculated as indicated in Figure 2 and explained in the methods section.

We followed by generating whole-transcript polyA^+^ RNA-seq datasets from the heads of all three species entrained at the 3 different temperatures. When examining the splicing pattern of *tim*, we could see that all species displayed dynamic changes in *tim-cold* and *tim-M* in response to temperature changes, with *tim-cold* increasing and *tim-M* decreasing at lower temperatures (Figure 3B and Figure S6). This suggests that these changes might be important for the behavioral changes shared by all species. However, we only detected *tim-sc* in *D.simulans* and *D. melanogaster*, which had very similar temperature-dependent alternative splicing profiles (Figure 3B and Figure S6). As mentioned above, this isoform appears only at 18°C, strongly suggesting that this isoform is responsible for the adaptation to cold temperatures, which is shared between *D.simulans* and *D.melanogaster*. The results displayed above showed a strong correlation among the different fly species between the changing on *tim* mRNA isoforms and the capacity to adapt to colder temperatures. We also looked at changes in *per* 3’ UTR splicing in the different species between temperatures. *D.virilis* displayed no changes in *per* splicing, while the changes in *D.Yakuba* were mild, in both cases the spliced isoform dominated (Figure 3B and Figure S6B).

### Cold temperature decreases TIM-L levels by two independent mechanisms

As previously described (Majercak et al., 1999), we found that the levels of TIM-L protein are significant downregulated at 18°C compared to 25°C or 29°C. We could not clearly detect or differentiate the other isoforms by Western Blot (data not shown). Nevertheless, TIM-COLD has been reported to be expressed by Boothroyd et al. (2007a). *tim-M* mRNA is very abundant but we could not see a protein of the predicted size in the western blotts, which strongly suggests that that this isoform is poorly and/or not translated. Hence, expression of this isoform could serve as a way to regulate the amount of *tim-L* production. This agrees with a recent publication that identified this mRNA isoform as non-coding (Shakhmantsir et al., 2018). We also could not univocally detect TIM-SC. While we observed bands of TIM-SC expected size in the TIM immunoblot, these bands are present at all temperatures (data not shown). We believe that these bands might represent canonical TIM degradation products of similar size than TIM-SC (see below). In addition, the utilized TIM antibody was raised against the whole protein and detects TIM-SC with low efficiency (see below).

Interestingly, the lower levels of TIM-L at 18°C are not due to changes in total *tim* mRNA, as observed in our datasets (Figure S4) and reported by others (Boothroyd et al., 2007b). This suggests that the lower TIM levels are due to post-transcriptional and/or post-translational regulation. miRNA-mediated repression is one of the most common mechanisms of post-transcriptional control. Moreover, recent work demonstrated that *tim* mRNA is regulated by miR-276 (Chen and Rosbash, 2016). We then decided to determine whether *tim* is regulated by miRNAs in a temperature dependent way. To do so, we tested the genome-wide association of mRNAs with the ARGONAUTE 1 (AGO1, the only miRNA-RISC effector protein in *Drosophila* (Förstemann, Horwich, Wee, Tomari, & Zamore, 2007) at 18°C, 25°C and 29°C. Briefly, we performed AGO1 immunoprecipitation (IP) from fly heads followed by hybridization to oligonucleotide microarrays. In these datasets, we can only assess overall *tim* levels (as these microarrays cannot distinguish between *tim-L, tim*-*cold* and *tim-M*). We first confirmed that the overall distribution of AGO1-associated mRNAs (and hence miRNA-mediated regulation) is similar at all three temperatures (Figure S7). Interestingly, we observe changes in the association of *tim* to AGO1 (Figure 4A). Binding of *tim* to AGO1 is very strong at 18°C while there is very little or no association at 25 or 29°C. This patter is *tim*-specific and, when examining other core circadian clock components, we could not see such temperature-driven AGO1 association changes. Some are bound at all temperatures (like *Clk* or *vri)* and some do not bind at all (like *cry, cyc* and *per*; Figure 4A). Importantly, the probes in the oligonucleotide microarray do not allow distinguishing between *tim-L, tim-cold* and *tim-M*. To determine whether the association of the different *tim* transcripts to the miRNA-effector machinery is temperature dependent, we performed isoform-specific qPCRs from newly generated AGO1-IP samples. While the mRNA encoding the transcription factor CBT (a known miRNA-regulated RNA (Kadener et al., 2009; Lerner et al., 2015)) was strongly bound to AGO1 in all the assayed temperatures, we discovered that the association of *tim-L/M, tim-cold* and *tim-M* to AGO1 was significantly higher at 18°C (Figure 4B).

**Figure 4.**
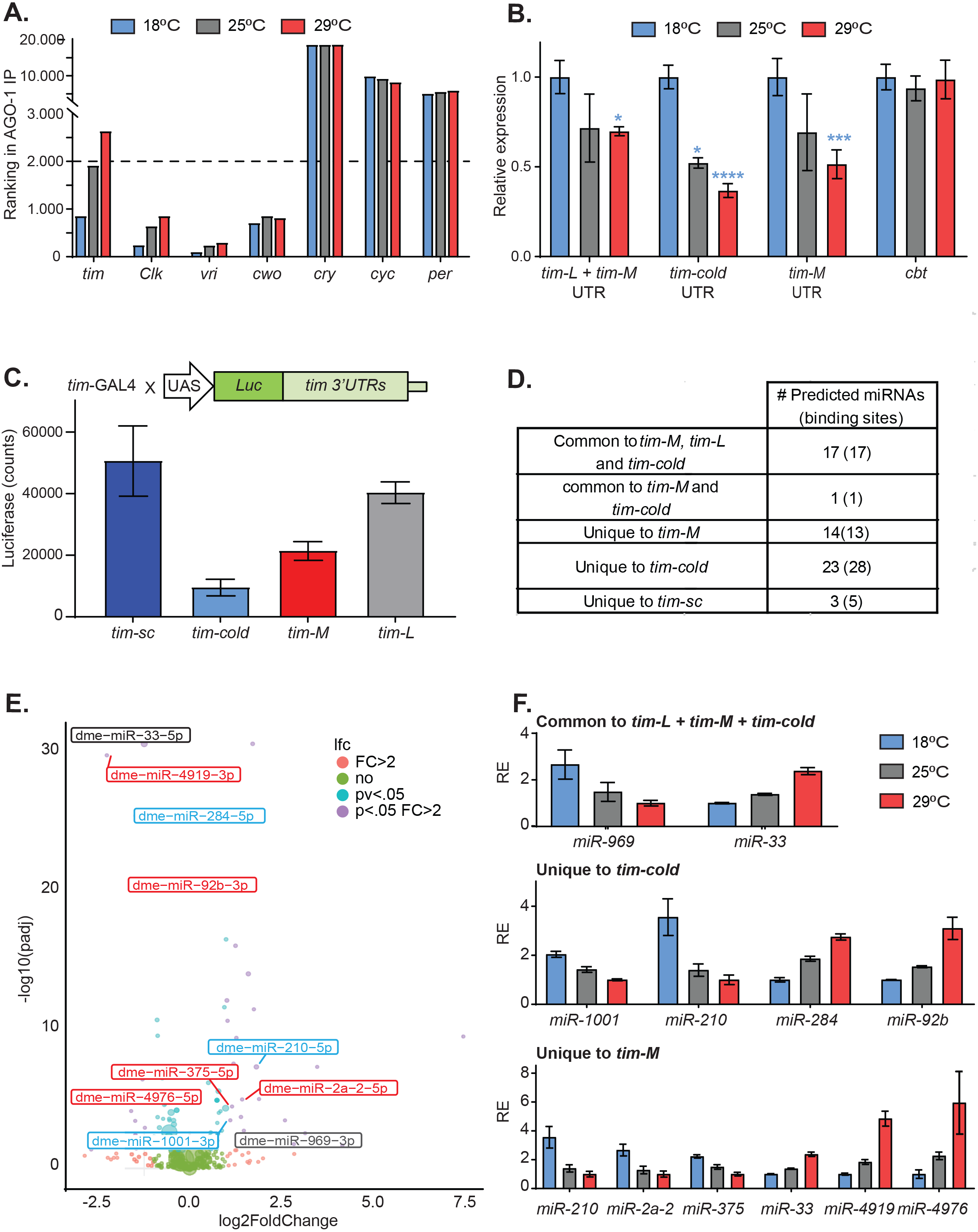
Alternatively spliced isoforms of *timeless* are post-transcriptionally regulated. **A.** *Tim* mRNA is bound to AGO-1 in a temperature dependent way. The graph represents the ranking of enrichment of RNAs in AGO-1 immunoprecipitates isolated from flies entrained at 18°C (blue), 25°C (grey) or 29°C (red). Briefly, we ranked RNAs according to their binding enrichment (lower ranking value corresponds to higher binding to AGO-1). To assess the ranking, we averaged the residuals in a linear model: signal in AGO1-immunoprecipitate/signal in input (n=2 for each temperature). **B.** *tim* isoform-specific qPCR of AGO1-bound mRNAs at 18°C (blue), 25°C (grey) or 29°C (red). Each bar represents an average of all time points normalized to a negative control (*cyc*). Blue starts represent statistical significance to 18°C of the indicated isoform and temperature determined performing multiple t-test analysis using the Holm-Sidak method. (*, pval<0.05; ***, pval<0.001; ****, pval<0.0001). Four different timepoints in each temperature were used as biological replicas. **C.** Luciferase assay of the wings of flies expressing *luciferase* fused to different *tim 3’UTRs*. We utilized the GAL4-UAS system to express the luciferase reporter. In all cases we utilized the *tim-gal* driver. All the UAS-Luciferase lines were inserted in the same genomic location using the Attb system. Blue and grey starts represent statistical significance to *tim-sc* or *tim-L*, respectively, determined by performing multiple t-test analysis using the Holm-Sidak method. (*, pval<0.05; ***, pval<0.001; ****, pval<0.0001. n=3). **D.** Summary table of the number of miRNAs binding identified in the different *tim* isoforms using TargetScan (Left column). **E.** Volcano plot showing differentially expressed miRNAs between 18°C and 29°C (n = 4 for each temperature). The color code represents the significance for each miRNA differentiating all the combinations for fold change > 2 (FC>2) and padj < 0.05. miRNAs with predicted binding sites present in any of the *tim* isoforms 3’UTR were identified using TargetScanFly.6.2 and are labelled in the graph (blue, grey or red for being putative targets for *tim-cold*, common sequence or *tim-M* 3’UTRs, respectively). miRNA abundance was assessed by small-RNAseq from AGO1 immunoprecipitates. **F.** Relative abundance of miRNAs that are differentially expressed (FC>2 and padj < 0.05) at different temperatures and that are predicted to target the 3’ UTR of the different isoforms. For each miRNA, the values have been normalized to the minimum and their relative expression (RE) is represented. miRNA abundance was assessed from small-RNAseq from AGO1 immunoprecipitations.

To determine the consequences of AGO1 binding on TIM expression, we generated flies carrying luciferase reporters fused to the different 3’ UTRs of *tim*. All transgenes were targeted and inserted in the same genomic location, and we expressed these UAS-transgenes using the *tim-gal4* driver. To minimize the effects of temperature on the GAL4 transgene, we performed the experiment at 25°C. The Luciferase levels of the transgenes carrying the 3’ UTRs of *tim-L* and *tim-sc* were high, suggesting little (or no) post-transcriptional control, at least at 25°C. Interestingly, the reporter carrying the 3’ UTR of *tim-cold* showed a strong decrease of Luciferase levels compared to the rest of the reporters, suggesting that the presence of this 3’ UTR strongly diminishes the translation potential of this isoform either by affecting the stability of the mRNA or, directly, by preventing its translation (Figure 4C). Similarly, the *tim-M* 3’ UTR reporter displayed lower luciferase levels, although the effect was milder than the one observed with *tim-cold* 3’ UTR.

To further understand the post-transcriptional regulation of *tim*, we analyzed the different 3’ UTRs for potential miRNA binding sites using Target Scan (Agarwal, Bell, Nam, & Bartel, 2015). Three of the isoforms, *tim-L, tim-cold* and *tim-M* contain a large number of miRNA binding sites, many of which are evolutionary conserved and highly expressed in fly heads (Table S3). *Tim-sc* contains only 5 miRNA binding sites and was not bound to AGO1 at any temperature demonstrating it is not regulated by miRNAs (Figure 4D).

The miRNAs responsible for the association of *tim* isoforms with AGO1 could target only a temperature-specific isoform, more than one isoform, be expressed in a temperature-dependent way or a combination of these. We therefore sequenced AGO1-associated miRNAs from heads of flies entrained to 12:12 LD cycles at 18, 25 and 29°C. While the expression of most miRNAs was not affected by temperature, a few of them were (Figure 4E, Table S4). We first focus on the miRNAs targeting the 3’ UTR region common to *tim-L, tim-cold* and *tim-M*. Of the 17 miRNAs predicted to target this region, only two change with temperature. One of them (miR-33) is modestly increased at 29°C in comparison with 18C. Notably miR-969, which contains a strong binding site in this 3’ UTR, is strongly (~3 fold) upregulated at 18°C, with intermediate expression at 25°C and low at 29°C (Figure 4F). Changes in the abundance of this miRNA might explain the temperature dependent association of *tim-L* to AGO1. However, *tim-M* and *tim-cold* contain additional 3’ UTR regions and these regions are predicted to be targeted by other miRNAs that are strongly regulated by temperature. For each of these *tim* isoform, half of the miRNAs predicted to bind them are highly expressed at 18°C and half highly expressed at 29°C (Figure 4F and Tables S3 and S4). This strongly suggests the existence of a mechanism for downregulating TIM-COLD and TIM-M at all temperatures. The fact that TIM-COLD is indeed present at 18°C (Boothroyd et al., 2007a; Wijnen et al., 2006), suggest that the main function of the miRNA-mediated regulation of this isoform is to impose a threshold for avoiding expression of this protein at temperatures other than 18°C.

In sum, these results, strongly suggest that, at 18°C, changes in the canonical TIM protein are due to the deviation of *timeless* transcription towards the production of a short isoform (*tim-sc*) and another isoform under strong post-transcriptional regulation (*tim-cold*). In addition, at this temperature even *tim-L* is subjected to a stronger post-transcriptional control by miRNAs. Last, *tim-M* is also regulated post-transcriptionally at all temperatures (similar to *tim-cold*).

### Overexpression of the different TIM isoforms alters circadian behavior differently

We next sought to determine the functionality of the different TIM isoforms. To do so we generated plasmids expressing the different TIM protein isoforms fused to a C-terminal FLAG tag and the same 3’ UTR under the control of the UAS-transgene (Figure S8A). To verify the expression of TIM from these plasmids we co-transfected each one of them into *Drosophila* S2 cells (which do not express *tim*) together with a GAL4 expressing plasmid. As expected, using an anti-FLAG antibody we observed the bands of predicted size for TIM-SC, TIM-M, TIM-COLD and TIM-L (Figure S8B left). Interestingly, anti-TIM antibodies can efficiently detect TIM-L and TIM-COLD but barely detected TIM-SC (Figure S8B right). Moreover, we observed a degradation product that overlaps with TIM-SC, demonstrating that it is very difficult (if not impossible) to detect this isoform univocally by western blot. We followed by generating flies containing these plasmids by targeting them into the same *attB* insertion site, in order to obtain flies that can express the proteins at similar levels. We then over-expressed each of these TIM protein isoforms using the *tim-gal4* driver and determined the locomotor activity patterns of these flies at 25°C. When kept under a 12:12 LD cycle, we observed a clear advance in the start of the evening activity peak in the *tim-sc* overexpression flies compared to the *tim*-Gal4 control (Figure 5A, B). This, together with the fact that in DD overexpression of TIM-SC led to a one hour shortening of the period compared to the control (Figure 5C), suggests a role for *tim-sc* mRNA and protein in the behavioral advance observed in *wild type* flies at 18°C.

**Figure 5.**
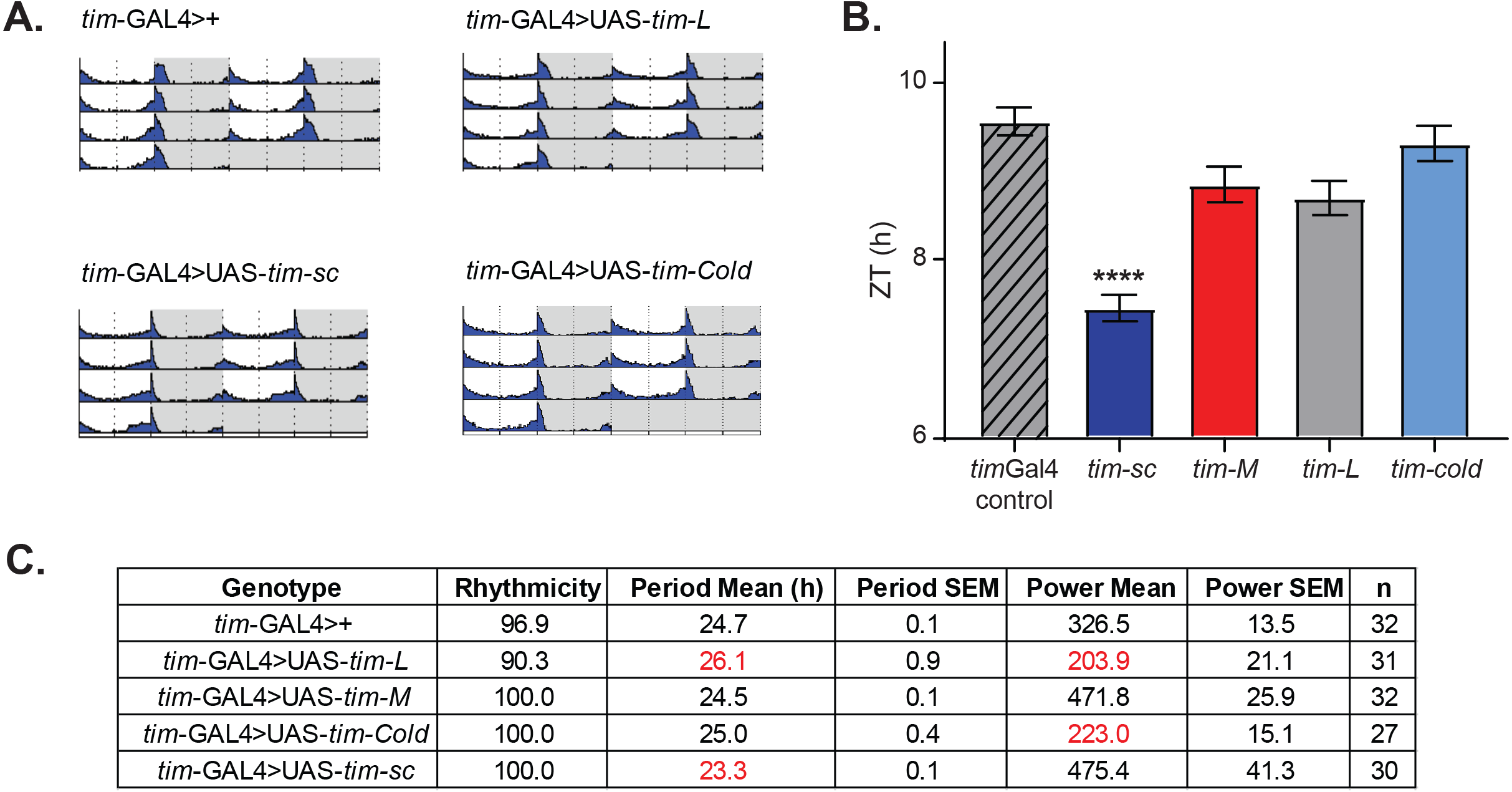
Over-expression of *tim* isoforms at 25°C results in various behavioral defects. **A.** Actograms of flies of the indicated genotypes in LD12:12 at 25°C. In all cases we over-expressed the indicated isoform utilizing the *tim-gal4* driver. Light phase is represented in white and dark-phase in grey. **B.** Quantification of the start of the evening peak in the last 3 days of LD. Stars represent statistical significance to *tim*-Gal4 control calculated by one-way ANOVA. (*, pval<0.05; **, pval<0.01 ***, pval<0.001; ****, pval<0.0001. n=27-31). **C.** Summary table of the behavior in DD indicating % of rhythmicity, period length, power and their SEM.

Additionally and as previously described by Yang and Sehgal (2001), overexpression of TIM-L at 25°C resulted in a high percentage of flies displaying long, weak or no rhythms in DD conditions (Figure 5C). Overexpression of TIM-COLD also resulted in weak and/or longer rhythms although at a lower degree than TIM-L, which suggests that the two proteins are not fully equivalent (although both are functional). Surprisingly, overexpression of TIM-M did not significantly change the locomotor activity pattern neither in LD nor in DD, suggesting that this protein is non-functional (Figure 5).

### Elimination of tim-sc results in changes in tim processing and locomotor activity

To definitively establish whether *tim-sc* is functional, we generated flies in which the cleavage and polyadenylation site used for the generation of *tim-sc* mRNA is mutated by CRISPR (40A flies, Figure 6A). As expected, these flies do not express *tim-sc* at any temperature (see qPCR in Figure 6B). We followed by assessing the locomotor behavior of 40A mutants and their isogenic controls. Control flies presented two different responses when transferred to 18°C: an advance of the evening peak as well as a decreased night activity in comparison to the flies maintained at 25°C (Figure 6C, 6D and 6E). In addition, the activity patterns of the 40A flies are strikingly similar at the different temperatures. At 18 and 25°C, 40A mutants display lower activity than control flies only during the night while at 29°C, these 40A flies were less active both in the light and dark periods (Figures 6D and S9). Even more importantly, and opposite to TIM-SC overexpression, 40A mutants display a significant delay in the time of evening activity onset specially at 18°C but also more mildly at 25°C (Figures 6C and 6E).

**Figure 6.**
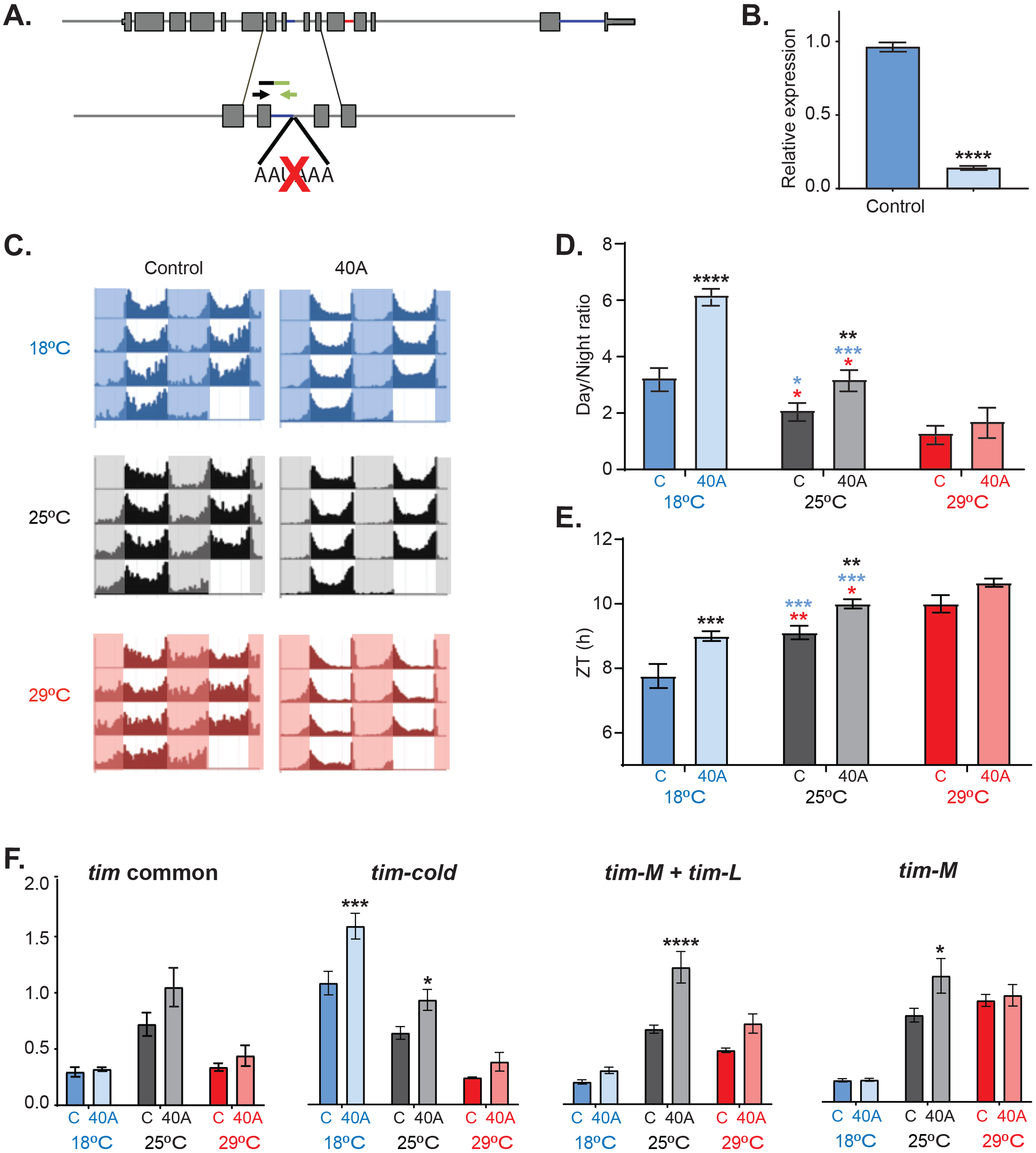
*Tim-sc* null flies present behavioral and molecular defects. **A.** Schematic representation of the mutation performed by CRISPR used to generate *tim-sc* null flies (40A flies). The black and green arrow represent the location of the forward and reverse primers used for the verification of *tim-sc* depletion. **B.** Levels of *tim-sc* after the genetic manipulation depicted in A. The displayed qPCR result was obtained using RNA obtained from heads of flies entrained at 18°C. (n=5). **C.** Average actograms of control (left) and 40A (right) flies at 18°C (blue), 25°C (grey) and 29°C (red) in LD12:12 cycles. (N=23-32). **D.** Ratio between total day and night-time activity at 18°C (blue), 25°C (grey) and 29°C (red) in 12:12LD. Blue and red stars represent statistical significance between flies kept at 18°C or 29°C and those at 25°C, while black ones represent differences to the control flies. Multiple t-test analysis using the Holm-Sidak method. (**, pval<0.01 ***, pval<0.001. n=16-31). **E.** Quantification of the start of the evening peak in day 3 of LD. Blue and red stars represent statistical significance between flies kept at 18°C or 29°C and those at 25°C, while black ones represent differences to the control flies. Multiple t-test analysis using the Holm-Sidak method. (**, pval<0.01 ***, pval<0.001. n=16-31). **F.** Assessment by qPCR of exons5-6 (common in all *tim* isoforms, left) *tim-cold* (middle left), *tim-L + tim-M* (middle right) and *tim-M* transcription (right) at 18°C (blue), 25°C (grey) and 29°C (red) at ZT15 in 12:12LD. Multiple t-test analysis using the Holm-Sidak method. (**, pval<0.01 ***, pval<0.001. n=5).

Finally, we measured the levels of the other *tim* isoforms in control and 40A flies at 18, 25 and 29°C in order to assess the molecular consequences of the disruption of *tim-s* production. *Tim-sc* mutant flies do not show changes in the overall levels of *tim* mRNA (as assessed by determining the levels of the constitutive exon 5 and 6; Figure 6F left). However, elimination of *tim-sc* results in an increase in the levels of *tim-cold* both 18 and 25°C at the maximum expression timepoint (ZT15; Figure 6E). *tim-L/M* levels also are increased in the mutant, but only at 25°C.

These results strongly suggest that both TIM-SC protein and *tim-sc* production can regulate the daily pattern of locomotor activity and the response to temperature changes. Importantly, we still observe some degree of temperature adaptation in 40A mutants, which we postulate are mediated by the increased production of *tim-cold*.

### Tim alternative splicing is regulated directly by temperature in-vitro and in-vivo

To get insights into the mechanism by which temperature regulates *tim* alternative splicing, we determined if *tim*^*01*^ and *per*^*01*^ mutants also change *tim* alternative splicing in a temperature-dependent way. Hence, we entrained control and both mutant lines in LD conditions at 18°C and at 29°C, collected flies every four hours, extracted RNA from fly heads from a mix of all time points and performed qPCR. We observed a similar trend in all three fly lines, namely: an increase in *tim-cold* and *tim-sc* and a decrease in *tim-M* at 18°C (Figure 7A). This demonstrates that, as previously shown for *per* temperature-dependent splicing, *tim* thermosensitive splicing events are independent of the circadian clock.

**Figure 7.**
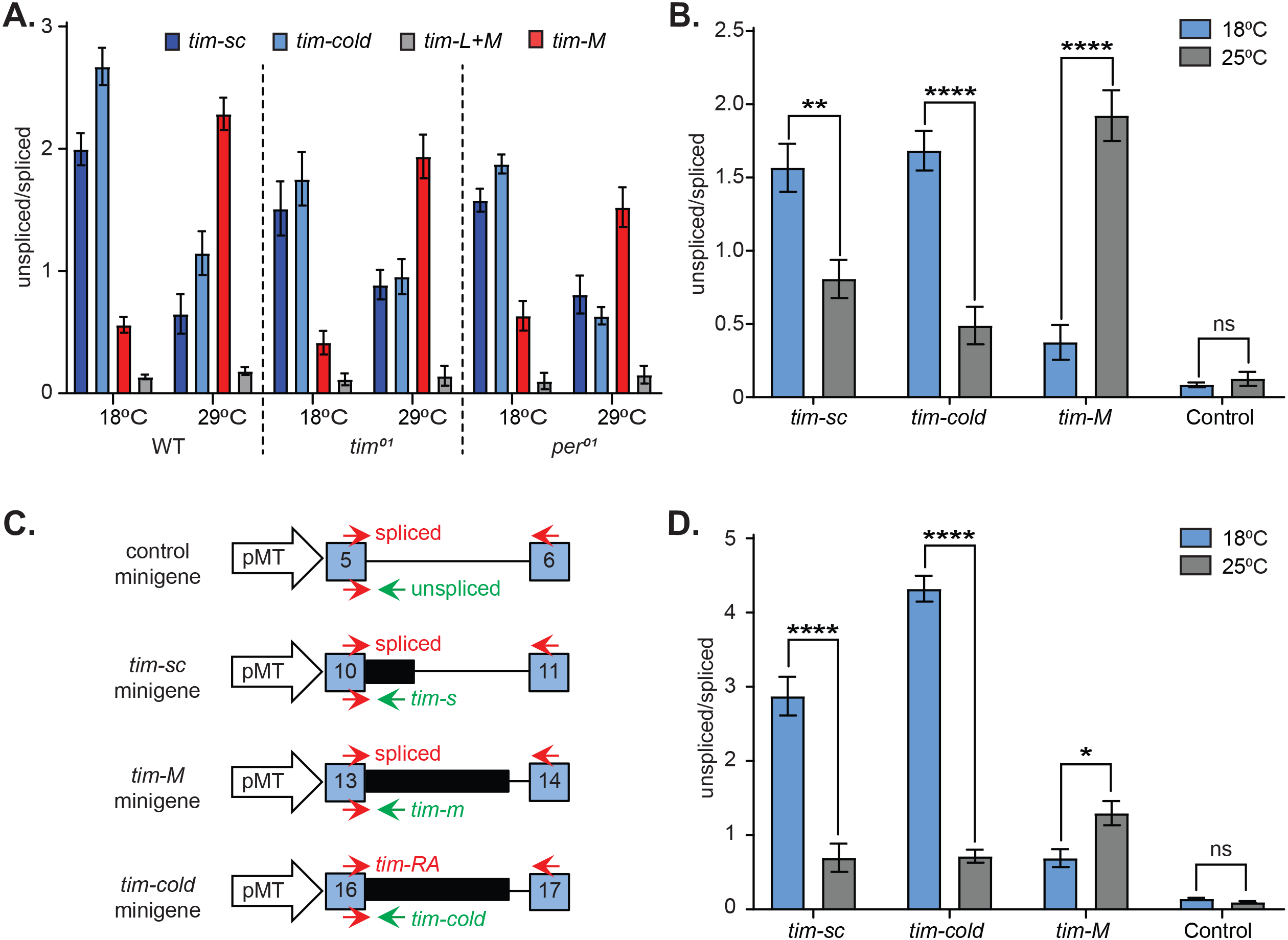
Splicing of *timeless* is temperature dependent and clock independent. **A.** qPCR results from entrained WT as well as *tim*^*01*^ and *per*^*01*^ clock mutant flies at 18°C or at 29°C. Each bar represents the ratio between the unspliced and splice variant of *tim-sc* (dark blue), *tim-cold* (light blue), *tim-m* (red) and *tim* control (grey). **B**. *Drosophila* S2 cells transfected with pMT-*Clk* display the same *tim* alternative splicing patterns observed in living flies at 18°C (blue) or at 25°C (grey). Each bar represents the ratio between the unspliced and splice variant of *tim-sc, tim-cold, tim-m* and *tim* control. Multiple t-test analysis using the Holm-Sidak method. (**, pval<0.01 ***, pval<0.001. n=3). **C.** Scheme of the different *tim* isoform minigenes utilized in (D) and location of the primers used to measure the spliced (top) and unspliced (bottom) of each intron. **D.** Unspliced/spliced ratios for the indicated minigenes at 18°C (blue) or at 25°C (grey). Multiple t-test analysis using the Holm-Sidak method. (**, pval<0.01 ***, pval<0.001. n=3).

The splicing sites flanking alternatively spliced exons are usually weaker and temperature is known, at least *in vitro*, to strongly influence the efficiency of splicing. Hence, we determined the strengths of the different splice sites in *tim* using a publicly available software (Reese et al., 1997). Almost all *tim* constitutive introns exhibited strong 5’ and 3’ splice site strengths (above 0.6 scores, Figure S10). Interestingly, the introns retained in *tim-M* and in *tim-cold* have strong 5’ splice sited and a weak 3’ splice site. This suggests that at least part of the mechanism underlying the production of these isoforms is due to the presence of weak 3’ splice sites. However, the splice sites of the intron associated with *tim-sc* bear strong 5’ and 3’ splice site (**Figure S10**), suggesting that the increase of *tim-short* at lower temperatures might be the result of a more efficient cleavage and polyadenylation rather than changes on the splicing efficiency. In addition, we observed that *D.melanogaster, D.simulans* and *D.Yakuba* possess strong 5’ splice sites and weak 3’ splice sites of the introns retained in *tim-cold* and *tim-M* whose alternative splicing is temperature sensitive in these species (Figure S10). Interestingly both *tim-cold* and *tim-M* respond in opposite ways to changes in temperatures (with *tim-cold* strongly retained at 18°C and *tim-M* more skipped at this temperature).

*Tim* was one of the few genes in which we observed temperature dependent changes in alternative splicing (data not shown). Hence, it is possible that *tim* alternative splicing *per* se could be thermosensitive. To test this possibility, we utilized the embryonic cell line Schneider 2 (S2 cells). These cells do not express most circadian components including *Clk, per* and *tim*. However, *tim* expression can be induced by expression of CLK (McDonald & Rosbash, 2001). So, we transfected these cells with a plasmid driving expression of *Clk*. We used an inducible vector in order to bypass any transfection variation that might arise due to different temperature conditions (Lerner et al., 2015). After the induction of *Clk*, we cultured the cells either at 25°C or 18°C for 24 hours. We then collected the cells, extracted RNA and characterized the splicing pattern of *tim* by qPCR. For each of the alternative isoforms, we measured the ratio between the unspliced and spliced variants and compared these ratios between the two temperatures. Indeed, we completely reproduce the results obtained *in* vivo, namely: *tim-cold* and *tim-sc* were higher at 18°C, while *tim-M* showed the opposite trend (Figure 7B). However, we did not observe any temperature-induced changes of a constitutive *tim* exon (Control, Figure 7B) demonstrating that this effect is specific for the thermosensitive introns.

One intriguing possibility is *tim* splicing *per se* functioning as a thermometer. This could be accomplished if temperature could, for example, expose binding sites for a specific splicing factor or if the splicing of these introns is temperature dependent. To test this possibility, we generated three different *tim* minigene reporters consisting of the exons and intron involved in each event of *tim* alternative splicing (Figure 7C). As observed *in vivo* and in the context of the whole *tim* gene in S2 cells, we observed a temperature-dependent effect on the splicing of the three different *tim* minigenes. We did not observe any temperature dependence in the splicing of a minigene generated from a constitutive exon (Figure 7D), demonstrating a direct effect of temperature on *tim* alternative splicing.

## DISCUSSION

In this study we show that temperature dramatically and specifically changes the splicing pattern of the core circadian component *timeless*. We found that the lower levels of the canonical TIM (TIM-L) protein at 18°C are due to the induction of two cold-specific splicing isoforms (*tim-cold* and *tim-short&cold*). *Tim-cold* encodes a protein very similar to TIM-L but it is under strong post-transcriptional control, as showed by AGO1-IP and *in vivo* luciferase reporters. Moreover, *tim-sc* encodes a short TIM isoform which results in an advanced phase of the circadian clock when overexpressed. Interestingly, these changes in *tim* splicing patterns are conserved across several *Drosophila* species and correlate well with the capacity of the species to adapt their activity to temperature changes. We then generated flies in which the production of *tim-sc* is abrogated. These flies display altered patterns of locomotor activity at 18°C and 25°C as well as altered expression of the remaining *tim* isoforms demonstrating the importance of *tim-sc* production. Last, we showed that the temperature-dependent changes in *tim* alternative splicing are independent of the circadian clock. Moreover, we could reproduce them in *Drosophila* S2 cells either promoting endogenous *tim* expression using the p-Act-*Clk* plasmid or utilizing splicing minigenes. The latter results strongly suggest that *tim* intronic sequences themselves are the temperature sensor for these splicing changes.

The fact that despite being the most abundant RNA isoform, we could not detect TIM-M suggests that this RNA is poorly (or not) translated. Alternatively, the protein could be quickly degraded. However, the protein is fairly stable as we could detect large amounts of TIM-M upon overexpression (Figure S8). Moreover, our luciferase reporter experiments suggest that little protein is produced from this RNA, likely by regulation by miRNAs. It is also possible that this RNA isoform is not even exported from the nucleus and nuclear/cytoplasmic fractionation experiments could be useful to test this possibility. Regarding *tim-cold*, this isoform is strongly regulated by mIRNAs at all temperatures and was reported (Montelli et al., 2015) to display weaker binding to CRY in yeast. Our results with the *tim-sc* null mutant provide support for a specific (and even dominant) function of TIM-COLD. We observed that *tim-sc* mutants have increased levels of *tim-cold* mRNA at 25°C. At this temperature these flies behave as if they were at lower temperature (low daytime activity and advanced evening peak of activity). Similar to *tim-cold, tim-sc* seems to work both at the RNA and protein levels. On one hand, the increase in the production of *tim-sc* at 18°C helps regulate the amount of *tim-L* and *tim-cold* produced at this temperature (Figure S8). In addition, overexpression of TIM-SC advances the phase and shortens the period of the circadian clock, suggesting that expression of this isoform might mediate some of the changes seen upon introduction of flies at 18°C. An accompanying manuscript (Foley et al., Submitted) shows that the splicing factor *psi* regulates *tim* alternative splicing in the same direction as when transferring the flies to colder temperatures. However, *psi* expression and activity are not modulated by temperature, suggesting that the temperature is sensed by a different system.

While *tim-cold* and *tim-M* are strongly post-transcriptionally regulated at all temperatures, the RISC binds also stronger to *tim-L* at 18°C. Indeed, *tim* has been reported to be regulated by miRNAs276a (Chen and Rosbash, 2016). Our miRNA-seq experiments suggest that miR-969 might be responsible for this temperature-specific regulation. However, it is possible that other miRNAs are involved and/or that other RNA binding proteins enhance the use of some miRNAs in a temperature-dependent way. CRISPR experiments targeting the miR-969 site could help distinguish between these possibilities. The miRNA profiling and AGO-IP experiments also suggest that other mRNAs might be regulated by miRNAs in a temperature-dependent way. It will be really interesting to match those datasets in order to understand how the transcriptional and post-transcriptional expression programs are regulated by temperature. Despite the large number of RNAs and miRNAs that are regulated in a temperature-dependent manner, the impact of temperature in alternative splicing seems to be quite restricted. Although we performed only a superficial analysis of the data, only *tim, per* and *Hsf* show evident temperature-dependent regulation of splicing (data not shown). Splicing is strongly temperature dependent, at least *in vitro*. Flies might have mechanisms that make the outcome of splicing temperature independent. This could be achieved by compensatory changes in chromatin structure, RNA pol II elongation rate or RNA editing which is known to be altered by temperature (Buchumenski et al., 2017). In addition, the specificity of the splicing strongly suggests that the changes in *tim* splicing are unlikely to be due to the expression or activation of a specific splicing factor. This possibility is supported by the findings reported in the accompanying manuscript by Foley et al. showing that while *psi* regulates *tim* splicing, this factor on its own cannot explain the temperature sensitivity of these alternative splicing events. Importantly, our results demonstrate that the introns in isolation are still able to sense and respond to the temperature changes (Figure 7). These results suggest that RNA structure likely plays a key role in making this splicing events temperature sensitive. For example, temperature could change the accessibility of *psi* binding sites in a temperature dependent fashion. Interestingly, each temperature-driven splicing event might operate differently. For example, production of *tim-sc* results from the competition between splicing and a cleavage and polyadenylation event. On the other hand, production of *tim-M* and *tim-cold* depends on an intron retention event. Interestingly, both introns have weak 3’ splice sites but display opposite temperature sensitivities, which suggest that their regulation depends on specific structures and/or sequences and are not the results of general effects of temperature on splicing.

We observed that flies that cannot generate *tim-sc* re-route *timeless* transcription towards the other isoforms, mainly *tim-cold* (at both 18 and 25°C). These mutant flies also display altered behavior. But what are the molecular actors responsible for these defects? *Tim-sc* mutant flies display higher levels of *tim-cold* and *tim-L* at 25°C. It is difficult to rationalize how increased levels of *tim-L* could trigger behavioral patterns observed in control flies at 18°C. We indeed favor the possibility that the behavior observed in these flies at 25°C is due to increased expression of *tim-cold* mRNA and protein. Previous work showed that TIM-COLD might act differently than TIM-L (Montelli et al., 2015). Importantly, the increase in TIM-COLD protein in the mutant flies might be significantly higher than the one on its RNA levels. This could happen if for example, the increase in *tim-L* mRNA indirectly increases TIM-COLD levels by titrating miR-969 or other miRNA targeting both to *tim-L* and *tim-cold* mRNAs. Additional CRISPR mutants (i.e. abrogating *tim-cold* expression or making it constitutive) could help to completely understand the mechanism behind this regulation.

Based on these and previous results we propose the following model in which miRNA-mediated control imposes temperature-dependent thresholds for protein expression for the different *tim* isoforms (Figure 8). At 25°C, *tim-L* and *tim-M* are produced. Both are moderately regulated by miRNAs, due to sequences present in their last exon. *Tim-cold* production is low at 25°C and the miRNA-mediated control is enough to abolish most (or all) protein expression from this isoform at this temperature. In addition, *tim-M* is target of many additional miRNAs, some of which are not differentially regulated by temperature, while others are higher at 29°C or 18°C. Therefore, we predict that no TIM-M is produced at any temperature, and the production of this isoform has a regulatory role (as recently suggested by Shakhmantsir et al. (2018)). When the temperature is decreased, the strong increase in *tim-cold* overcomes miRNA-mediated repression and some TIM-COLD is produced. In addition, *tim-sc* RNA and protein are produced. We hypothesize that both TIM-COLD and TIM-SC contribute to the phase advance and lower night activity observed at 18°C. TIM-SC lacks the cytoplasmic retention domain and increased levels of this protein might lead to the phase advance observed at 18°C. At 29°C, *Tim-M* is increased but TIM amounts are maintained constant likely due to higher production rates of TIM (due to a more modest association of *tim-L* with AGO1). In this way, *tim* co and post-transcriptional regulation can modulate the amounts and type of TIM proteins to facilitate temperature adaptation. It is possible that these changes in alternative splicing could regulate other situations in which the circadian clock requires adjustments of TIM levels and/or activity independently of CLK-driven transcription. These could include entrainment by temperature or light as well as cell to cell synchronization. Moreover, as *tim* is being alternatively spliced, it is even possible that some of these protein isoforms could be used by the cell as temperature sensors.

**Figure 8.**
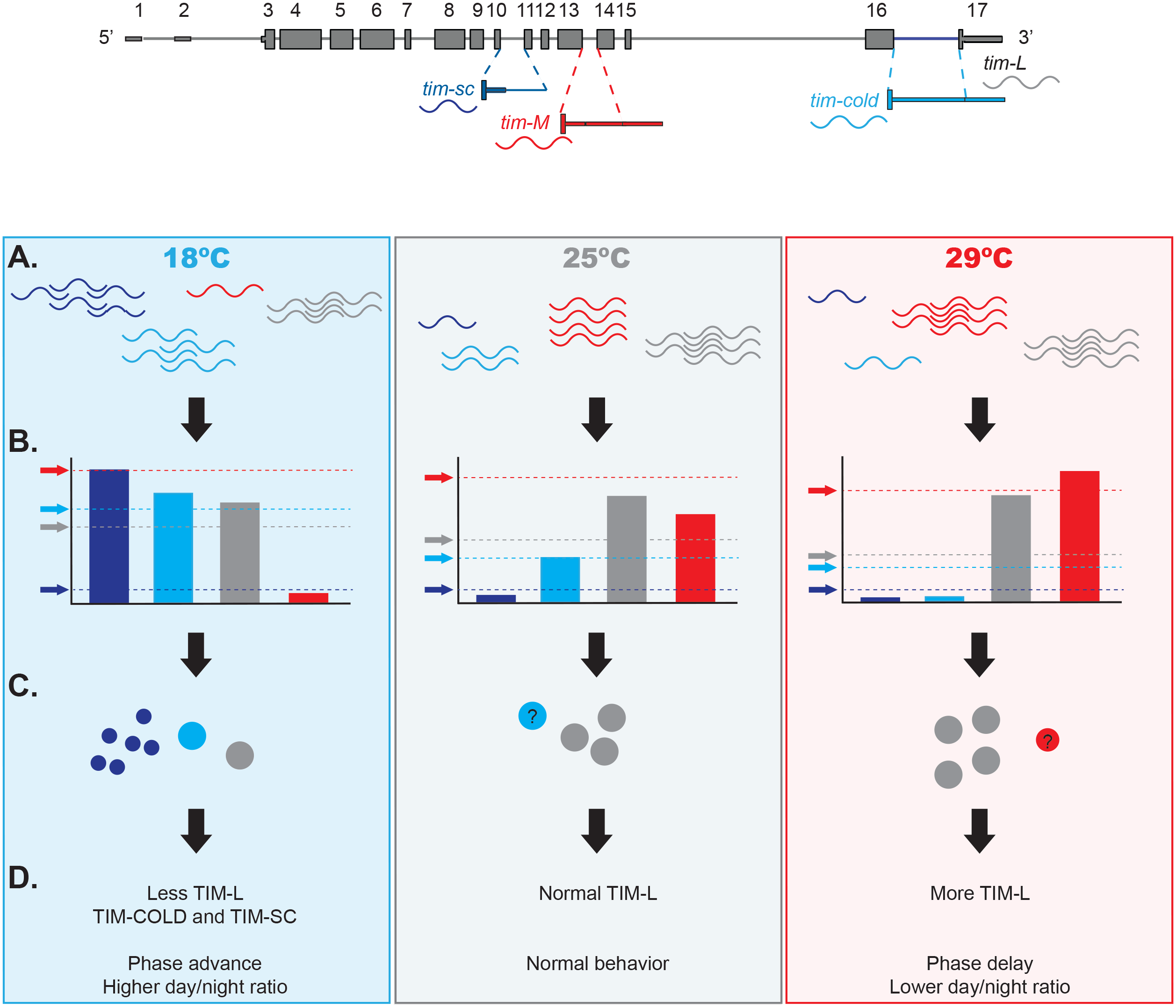
Model of the effects of thermosensitive alternative splicing of *timeless* in sensing and mediating temperature adaptation. **A.** Temperature affects differently each one of the four *tim* isoforms (*tim-sc, tim-cold, tim-M* and *tim-L*, represented in dark and light blue, red and grey respectively). While *tim-sc* and *tim-cold* are higher in colder temperatures, *tim-M* is only expressed at the highest temperatures. Nevertheless, transcription levels of the canonical isoform, *tim-L*, do not change. **B.** Temperature regulates expression of miRNAs, which set a threshold to the amount of mRNA that gets translated from each isoform. *tim-sc* has the lowest and *tim-M* the highest thresholds in all temperatures. *tim-cold*’s has the biggest temperature-dependent variation. **C.** These differences between the transcript levels and their thresholds results in changes in TIM expression as well as **D.** behavioral changes in the phase and day/night ratios that are linked to temperature adaptation.

In sum, in this study we determine that *tim* is extensively regulated by alternative splicing in a temperature dependent way. Moreover, we show that this regulation is, at least partially, responsible for temperature adaptation. We propose that this complex regulation of alternative splicing and miRNA-mediated control provides a mechanism for altering the relationship between TIM and PER proteins. This would allow the modulation of the circadian phase without affecting the period (which is strongly dependent on CLK and PER protein levels and activity). Last but not least, our data suggest that *tim* alternative splicing might act as a thermometer for the cell and might regulate temperature responses not related to the circadian clock.

## ACKNOWLEDGMENTS

We thank Avigayel Rabin and Reut Ashwall-Fluss for help with the alignment of the RNA seq data. This work was funded by the Binational Foundation (BSF Transformative Program), the ISF Legacy Program (Morasha, #1649/16) and the iCoreI-CORE Program of the Planning and Budgeting Committee in Israel (1796/12). All the RNA-seq data has been submitted to GEO (entries GSE124134, 124135, 124141, 123142, 124200 and 124201).

## AUTHOR CONTRIBUTIONS

NE generated the plasmids, performed the cell culture experiments and generate the CRISPR flies. AMA performed the behavioral analysis as well as the qRT-PCRs, analyzed the data, generated the figures and wrote the manuscript. OB generated the RNA-seq libraries, AGOIP experiments and western blots. ILP performed the computational analysis. RW performed the behavioral experiments in the non-melanogaster *Drosophila* species. SK designed the experiments and wrote the manuscript.

## DECLARATION OF INTEREST

The authors declare no competing interests.

## Materials and Methods

### Fly husbandry

*Drosophila yakuba, Drosophila simulans* and *Drosophila virilis* were obtained from the Drosophila Species Stock Center (DSSC) at the University of California, San Diego. CantonS flies were used as wild type strain for *Drosophila melanogaster*. Additionally, *tim*^*0*^ and *per*^*01*^ strains have been described previously in (Konopka and Benzer, 1971; Sehgal et al., 1994). Transgenic lines for the overexpression of different FLAG-tagged *tim* isoforms or luciferase fused to their 3’UTRs were generated by injecting the plasmids (described below) in a site-specific manner into the pattP2 site using the PhiC31 integrase-mediated transgenesis system (Best Gene Drosophila Embryo Injection Services). These transgenic flies were crossed to *tim-Gal4* driver (Blau and Young, 1999; Kaneko and Hall, 2000).

All crosses were performed and raised at 25°C. Newborn adults were either maintained at 25°C or switched to colder (18°C) or warmer (29°C) temperatures as described in the text.

### Generation of *tim-sc* polyadenylation and cleavage signal mutants by CRISPR

The *tim-sc* mutant flies were generated following the protocol from Ge et al. (2016) with some modifications. pCFD5 plasmid (Addgene, plasmid #73914) was modified to exclude *Vermilion* and *attB* (pCFD5d). 3 gRNAs (one targeting *w* gene and two, *tim*) were generated from PCR templates and cloned intro pCFD5d (pCFD5d*w/tim*-1,2) as described in (Fu et al., 2014; Port and Bullock, 2016; Port et al., 2014). The donor template for homologous recombination contained a point mutation in an intronic sequence as well as a silent point mutation (one in each site targeted by the gRNAs). This fragment was then cloned into pUC57-*white*[coffee] (Addgene, plasmid #84006) between the SacI and HindIII sites (pUC57-*timShort*).

pCFD5d*w/tim*-1,2, pUC57-*white*[coffee] and pUC57-*timShort* were injected into *vas-Cas9 (y*^*1*^, *M(Ferguson et al.)ZH-2A*) flies (Ge et al., 2016) by Rainbow Transgenic Flies, Inc. (Camarillo, CA). G0 flies were crossed to second chromosome balancers and individual G1 *Cyo* flies from non-red-eyed populations were again mated to the 2^nd^ chr balancers. Individual G1 flies were subsequently genotyped to verify the deletion and the possibility of random integration either in the genome or at CAS9 cutting sites of the plasmid was assessed as described in (Ge et al., 2016). G2 *cyo* male and female flies from positive crosses were crossed to obtain non *cyo* homozygous stocks. Finally, the entire *tim* locus was sequenced in the *tim-sc* mutants and their isogenic controls to verify that there were no additional mutations in the *tim* locus.

### Plasmids for S2 cells transfection and/or injection into *Drosophila*

pMT-*Clk* was previously described in (Weiss et al., 2014) and p-Act-Gal4 (Addgene, plasmid #24344).

#### C-terminus FLAG-tagged UAS-*tim* overexpression plasmids

cDNA from Canton-S heads of flies reared at 18 (for *tim-sc* and *tim-cold*), 25 (*tim-L*) or 29°C (*tim-M*) were used to amplify each isoform. NotI and KpnI restriction sites and a FLAG-tag sequence (at the 3’ of the coding DNA but before the stop codon) were added by PCR. Each isoform was finally cloned into the pUASTattB vector between the NotI and KpnI sites.

#### UAS-*luciferase*-*3’UTR* of each *tim* isoform plasmids

A PCR product containing the *Firefly Luciferase* gene was cloned into pUASTattB (pUAST-*luc*-attB). The 3’ UTRs of all four isoforms were amplified from fly head cDNA and cloned into the pUAST-*luc*-attB vector between NotI and XhoI sites.

#### *tim* exon-intron-exon minigenes

A plasmid containing exon5-intron5-exon6 (*tim-control*), exon10-intron10-exon11 (*tim-sc*), exon13-intron13-exon14 (*tim-M*) and exon16-intron16-exon17 (*tim-cold*) fragments defined by KpnI and XhoI restriction sites was ordered from Syntezza Bioscience Ltd. This plasmid was then cut with KpnI and XhoI. The four fragments were extracted and cloned into pMT-V5 (Invitrogen).

### Cell culture and transfections

*Drosophila* Schneider-2 (S2) cells were cultured in 10% fetal bovine serum (Invitrogen) insect tissue culture medium (Biological industries) as in (Weiss et al., 2014). The experiments were performed at 18°C or 25°C, as described in the text.

### Luciferase activity assay

Luciferase activity from S2 cells was measured using the Dual Luciferase Assay Kit (Promega) following the manufacturer’s instructions as in (Krishnan et al., 2001; Weiss et al., 2014). Luciferase measurements from fly wings was performed as previously described (Plautz and Kay, 1999).

### RNA libraries preparation for RNA-seq

#### DGE (3’end-seq)

Flies were entrained for at least 3 days at 18°C, 25°C or 29°C and collected at 6 different timepoints (ZT3, ZT7, ZT11, ZT15, ZT19 and ZT23). RNA from the fly heads was extracted using TRIzol reagent (SIGMA). 3’-seq libraries were prepared as previously described (Shishki et al., Afik et al., 2017). These data have been deposited at NCBI GEO as XXX.

#### Total RNA-seq

RNA from fly heads collected at two or three different timepoints (ZT 3, 15 and 21 for 18C and 29C and ZT3 and 15 for 25C) was extracted using Trizol and used to generate polyA+ RNA-seq libraries. The library preparation procedure was modified from (Engreitz et al., 2013) as follows: 0.5μg of total RNA was polyA+ selected (using Oligo(dT) beads, Invitrogen), fragmented in FastAP buffer (Thermo Scientific) for 3min at 940 C, then dephosphorylated with FastAP, cleaned (using SPRI beads, Agencourt) and ligated to a linker1 (5Phos/AXXXXXXXXAGATCGGAAGAGCGTCGTGTAG/3ddC/, where XXXXXXXX is an internal barcode specific for each sample), using T4 RNA ligase I (NEB). Ligated RNA was cleaned-up with Silane beads (Dynabeads MyOne, Life Technologies) and pooled into a single tube. RT was then performed for the pooled sample, with a specific primer (5′-CCTACACGACGCTCTTCC-3′) using AffinityScript Multiple Temperature cDNA Synthesis Kit (Agilent Technologies). Then, RNA-DNA hybrids were degraded by incubating the RT mixture with 10% 1M NaOH (e.g. 2ul to 20ul of RT mixture) at 700 C for 12 minutes. pH was then normalized by addition of corresponding amount of 0.5M AcOH (e.g. 4ul for 22 ul of NaOH+RT mixture). The reaction mixture was cleaned up using Silane beads and second ligation was performed, where 3’end of cDNA was ligated to linker2 (5Phos/AGATCGGAAGAGCACACGTCTG/3ddC/) using T4 RNA ligase I. The sequences of linker1 and linker2 are partially complementary to the standard Illumina read1 and read2/barcode adapters, respectively. Reaction Mixture was cleaned up (Silane beads) and PCR enrichment was set up using enrichment primers 1 and 2: (5’AATGATACGGCGACCACCGAGATCTACACTCTTTCCCTACACGACGCTCTTCCGA TCT-3’, 5’-CAAGCAGAAGACGGCATACGAGATXXXXXXXXGTGACTGGAGTTCAGAC GTGTGCTCTTCCGATCT-3’, where XXXXXXX is barcode sequence) and Phusion HF MasterMix (NEB). 12 cycles of enrichment were performed. Libraries were cleaned with 0.7X volume of SPRI beads. Libraries were characterization by Tapestation. RNA was sequenced as paired-end samples, in a NextSeq 500 sequencer (Illumina).

### AGO1-IP followed by small RNA sequencing

The samples were collected at 18°C, 25°C or 29°C at four different timepoints (ZT3, ZT9, ZT15 and ZT21). AGO1 immunoprecipitation was performed as previously described (Kadener et al., 2009; Lerner et al., 2015). The small RNA libraries were constructed using NEBNext^®^ Small RNA Laibrary Prep Set for Illumina.

### AGO1-IP followed by Oligonucleotide microarrays

The samples were collected at 18°C, 25°C or 29°C. AGO1 immunoprecipitation was performed as previously described (Kadener et al., 2009; Lerner et al., 2015). INPUT and AGO IP Total RNA was extracted from fly heads using TRI reagent (Sigma) according to manufacturer’s protocol. cDNA synthesis was carried out as described in the Expression Analysis Technical Manual (Affymetrix). The cRNA reactions were carried out using the IVT kit (Affymetrix). Affymetrix high-density arrays for Drosophila melanogaster version 2.0 were probed, hybridized, stained, and washed according to the manufacturer’s protocol.

### Microarray Analysis

The R Bioconductor affy package (http://www.bioconductor.org) was used to normalize and calculate summary values from Affymetrix CEL files using gcRMA (Bioconductor). Gene ontology enrichment analysis was performed as described in Mezan et al. (2013). To determine AGO-IP enrichment ranking, we ranked the residuals from a linear model was between the normalized reads in the IP and the INPUT.

### RNA seq analysis

The total RNA sequencing reads were aligned to the *Drosophila melanogaster* genome version dm3. For the different species, alternative splicing proportion was calculated manually searching the exon-exon junction.

For the small RNA sequencing, the reads were processed using miRExpress pipeline (Wang et al., 2009) using miRBase 21 version. The different circadian timepoints were considered independent replicates (n=4) and the differential gene expression analysis was done using a negative binomial model using DeSeq2 package on R. A miRNA with p adjusted value less than 0.05 and an absolute log2(fold change) more than 1 was considered differentially expressed.

For the DGE data, the circadian analysis was performed using the package MetaCycle (Wu et al., 2016). For each temperature and circadian timepoint, two replicates were analyzed (n=2). To normalize over different library preparation, after normalizing by library size the counts were divided by the maximum in each replicate. Genes with more than two cero counts in any timepoint was discarded from further analysis. The amplitude for each replicate was then calculated as the maximum divided the minimum for each gene. JTK algorithm was used for the circadian analysis. A gene was considered as cycling if the JTK pvalue was less than 0.05 and the amplitude was more than 1.5. For these genes, the phase shift was calculated as the phase at 25°C minus the phase at 18ºC or 29ºC.

### TargetScan analysis of putative miRNA binding to each *tim* 3’UTR

As not all of the 3’UTRs for the different *tim* isoforms are annotated, TargetScanFly version 6.1 was run locally (which allows manual entry of the sequences of interest). UTR sequences were downloaded from UCSC dm6 27way conservation. This way we identified several putative miRNAs binding to each of the isoforms. We continued by manually looking at the abundance of this miRNAs (from the AGO1-IP followed by smallRNA sequencing) in order to determine which of those putative miRNAs are being expressed in the fly heads. Additionally, we defined as miRNAs that change in a temperature-dependent manner those miRNAs that had more than 2-fold change between 18 and 29°C and in which this difference was statistically significant (pval<0.05).

### Chromatin-bound (nascent) RNA

Performed as described in (Lerner et al., 2015).

### Real Time PCR analysis

Total RNA was extracted from adult fly heads (or brains) at the mentioned timepoints using TRI Reagent (Sigma) and treated with DNase I (NEB) following the manufacturer’s protocol. cDNA was synthesized from this RNA (using iScript and oligodT primers, Bio-Rad) and diluted 1:60 prior to performing the quantitative real-time PCR using SYBR green (Bio-Rad) in a C1000 Thermal Cycler Bio-Rad. Primers used for amplifying each isoform were: *tim-*sc (5’- AACACAACCAGGAGCATAC −3’ and 5’- ATGGTCCACAAATGTTAAAA −3’), *tim-*M (5’- GGAGACAATGTACGGACTC −3’ and 5’- ATTTCACACAGAGAGAGAGC −3’), *tim-*cold (5’- GCATCTGTGTACGAAAAGGA −3’ and 5’- ATGTAACCTATGTGCGACTC −3’), *tim-L* and *tim-*M (5’- CTCCATGAAGTCCTCGTTC −3’ and 5’- TGTCGCTGTTTAATTCCTTC −3’) and junction between exons 5−6 which are present in all isoforms (5’- AAAAGCAGCCTCATCAACAT −3’ and 5’- AGATAGCTGTAACCCTTGAG −3’).

The PCR mixture was subjected to 95 °C for 3 min, followed by 40 cycles of 95 °C for 10 s, 55 °C for 10 s and 72 °C for 30 s followed by a melting curve analysis. Fluorescence intensities were plotted versus the number of cycles by using an algorithm provided by the manufacturer (CFX Maestro Software, Bio-Rad). The results were normalized against *rp49* (5’- TACAGGCCCAAGATCGTGAA −3’ and 5’- CCATTTGTGCGACAGCTTAG −3’) and *tub* (5’- TGCTCACGAAAAGCTCTCCT −3’ and 5’- CACACACGCACTATCAGCAA −3’) levels.

### Western blotting

Protein was extracted from fly heads and WB performed as in (Weiss et al., 2014). Antibodies used: rat anti-TIM (1:30,000, a kind gift from Michael Rosbash), mouse anti-FLAG-M2 (1:20,000; Sigma F3165) and mouse anti-Tub (1:20,000, DM1A, SIGMA). Quantifications were done utilizing Image J software.

### Assessment of the locomotor behavior

Male flies were placed into a glass tube containing 2% agarose and 5% sucrose food and their activity was monitored using Trikinetics *Drosophila* Activity Monitors (Waltham, MA, USA). Flies were maintained for 4 days in 12:12 Light: Dark cycles (LD) and 5 days in constant darkness (DD). The experiments were performed at 18, 25 or 29°C, as indicated. Analyses were performed with a signal processing toolbox (Levine et al., 2002).

All the activity assessments were done in LD. For the calculation of the beginning of the evening peak, the activity of flies were analyzed individually on 30-minutes intervals. The criteria were: (1) have a trend of increasing activity, (2) no more than 1/3 of the bins can be outliers and (3) a reduction of less than 10% of the activity was not counted as an outlier. Only flies in which an evening peak was clearly distinguishable were included in this analysis.

---

**Figure S1.**
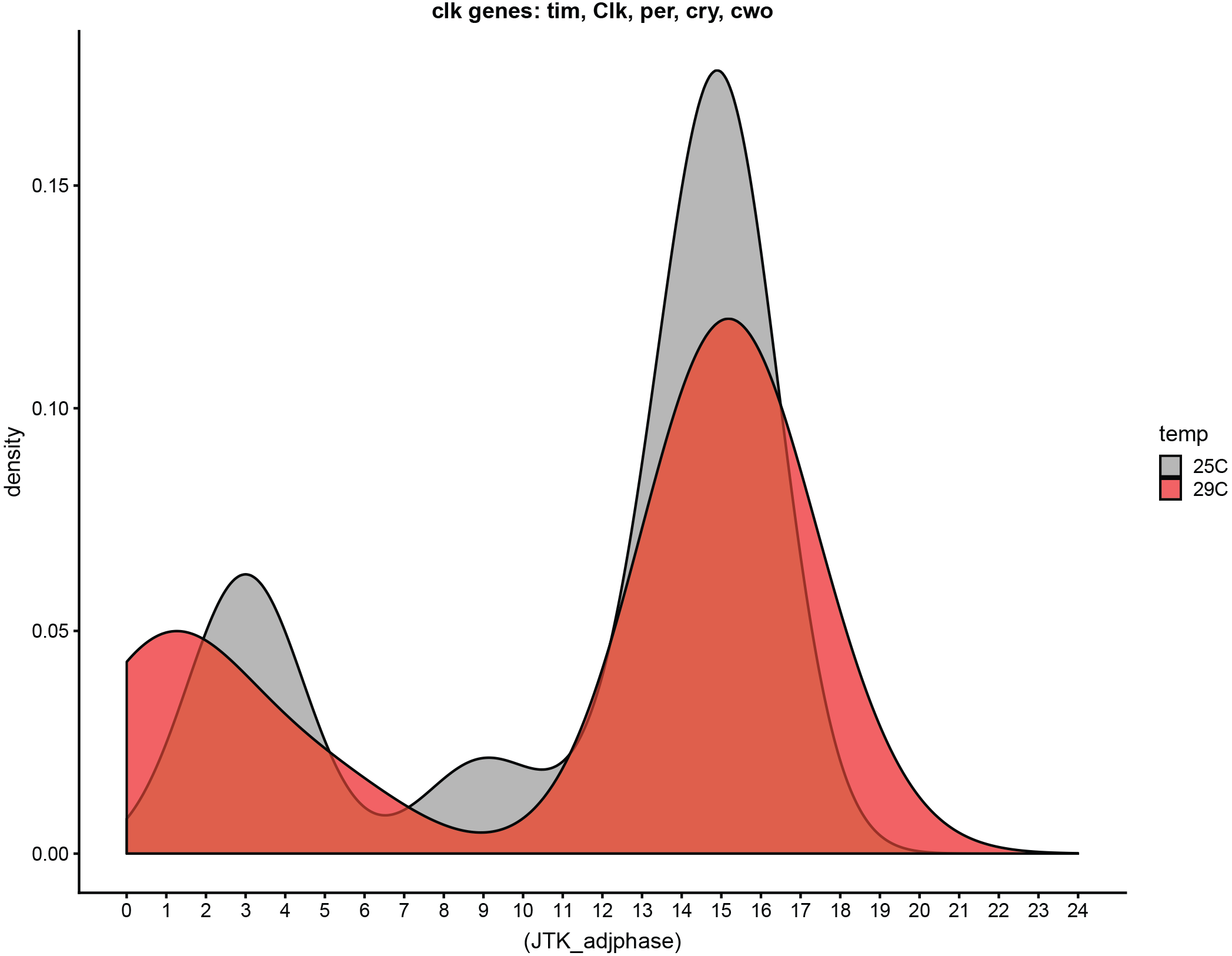
Density plot showing the phases of core clock genes cycling at both 29°C and 25°C. The red and grey shades indicate the phase of these genes at 29°C and 25°C respectively.

**Figure S2.**
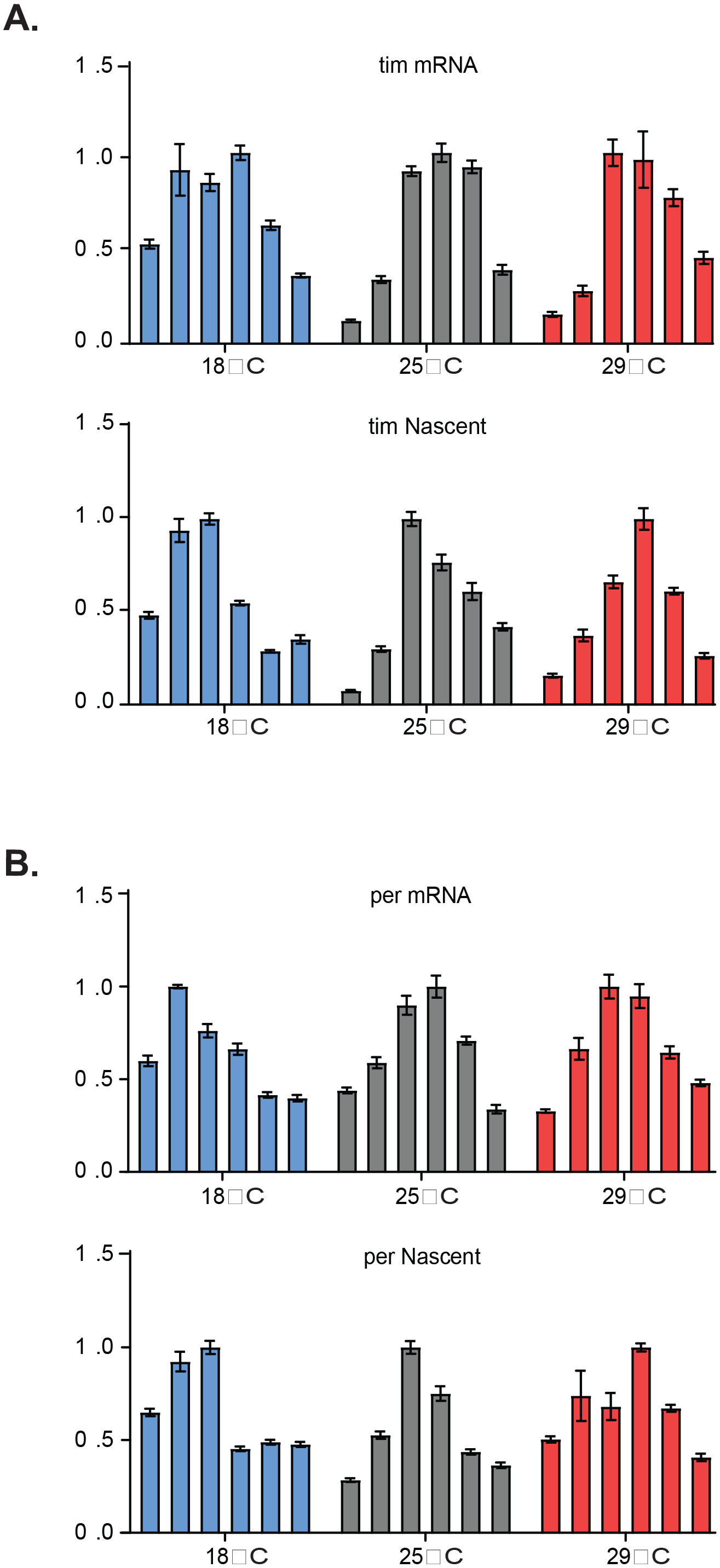
*Tim* transcription is advanced at 18°C compared to 25°C and 29°C. **A.** q-RT PCR results for *tim* mRNA, top, or chromatin-bound (nascent RNA) *tim*, bottom, levels on entrained WT fly heads at ZT3, ZT7, ZT11, ZT15, ZT19 and ZT23 (consecutive bars) at 18°C (blue), 25°C (grey) or 29°C (red). **B.** Same as in (A) but for *per*.

**Figure S3.**
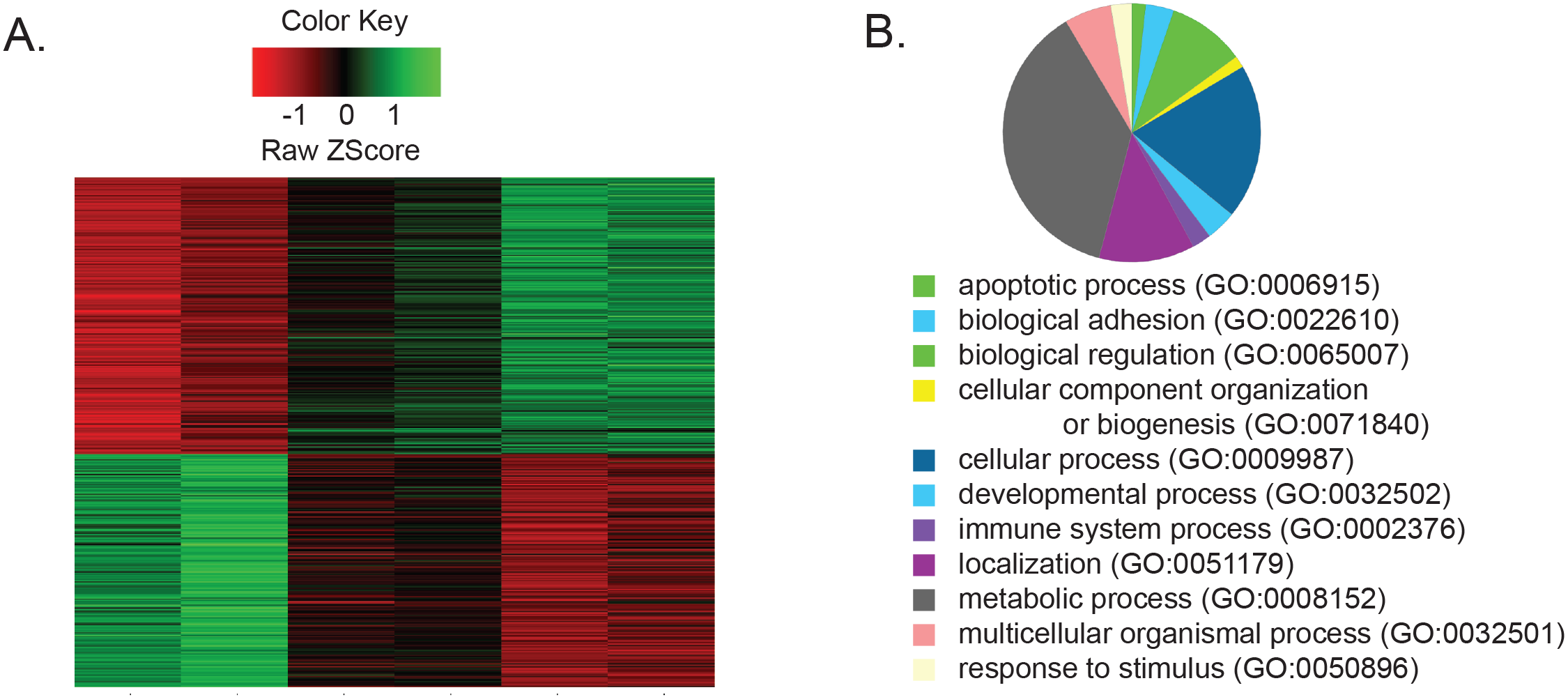
Temperature regulates gene expression. **A.** Heat map showing an unsupervised clustering of differentially expressed genes from a Microarray experiment performed on RNA extracted form entrained fly heads at three temperatures. Each lane represents a mix of all time points at a given temperature. **B.** A pie chart demonstrating enriched GO terms of the differentially expressed genes.

**Figure S4.**
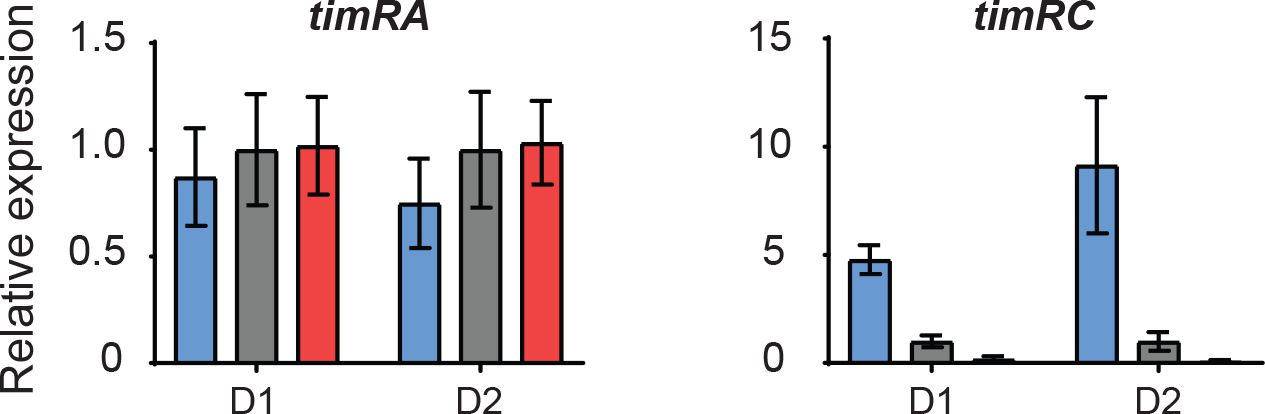
Relative expression of *timRA* and *timRC* in the two DGE described in Figure1. ***ti****m-RA* reports expression levels for *tim-L, tim-cold* and *tim-M*, as they share the 3’ end. For Each DGE dataset, we averaged the expression of the different *tim* isoforms over the day at each temperature. Results are shown normalized to 25°C.

**Figure S5.**
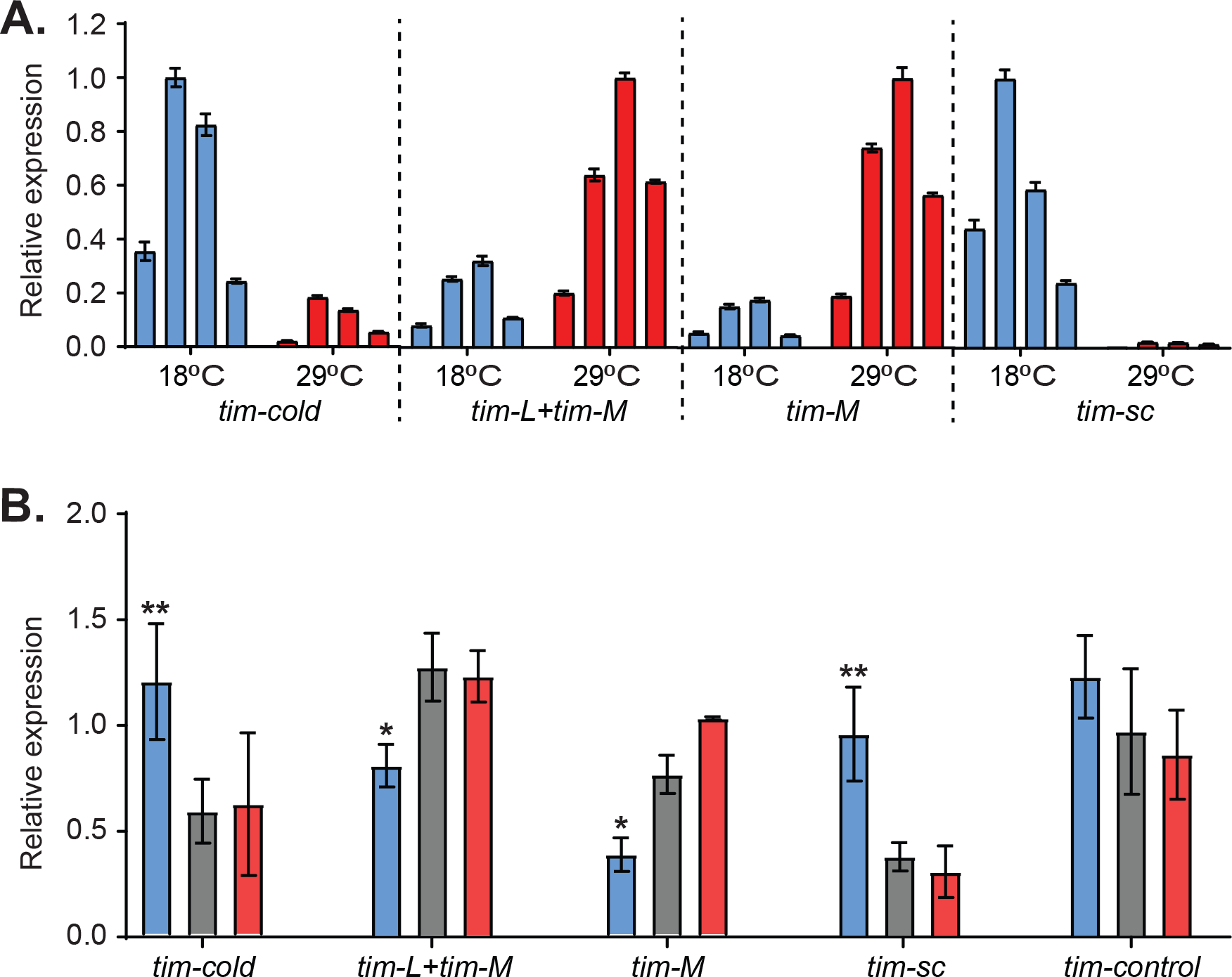
*Tim* expression is regulated by temperature co and post-transcriptionalyy. **A.** Isoform-specific q-PCR of *timeless* in the heads of flies entrained at 18°C (blue) and 29°C (red). Each bar represents a different time point (ZT3, ZT9, ZT15 and ZT21). Values have been normalized to the maximum of both temperatures. **B.** Quantification of the different *tim* isoforms in the adult brain at ZT11 after entraining the flies at 18°C (blue), 25°C (grey) and 29°C (red). Black stars represent statistical significance between flies kept at 18°C and those at 25°C. Multiple t-test analysis using the Holm-Sidak method. **, pval<0.01 ***, pval<0.001. n=16−31. n=3.

**Figure S6.**
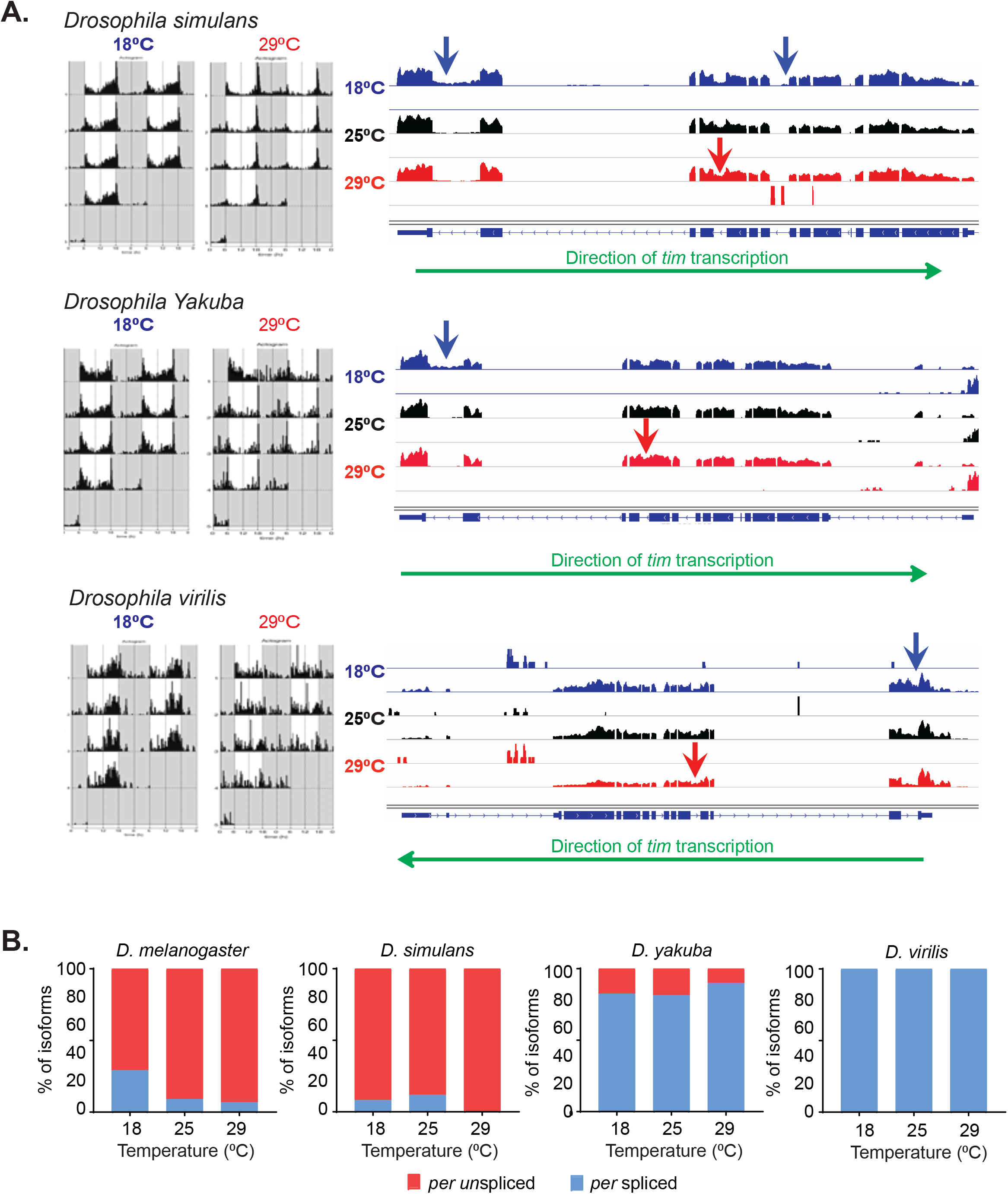
Temperature-specific alternative splicing is conserved across different *Drosophila* species. **A.** IGV snapshot of whole-transcript RNA-seq showing the *tim* locus in three additional species (*D. simulans, D. Yakuba* and *D. virilis*, top to bottom) at different temperatures. The upper panel (blue) corresponds to 18°C, the middle one (black) to 25°C and the lower one to 29°C. Arrows indicate the different areas unique to the different isoforms: *tim-sc* (light blue), *tim-M* (red) and *tim-cold* (blue). Green arrow below the IGV snapshots represent the direction of *tim* transcription. **B.** Quantification of the relative amount of *per-*spliced (light blue), *per-*unspliced (red) and *tim-L* (grey) from whole transcriptome (polyA^+^) mRNA-seq in all species at the indicated temperatures. The ratios between the isoforms were calculated as indicated in Figure 2 and explained in the methods section.

**Figure S7.**
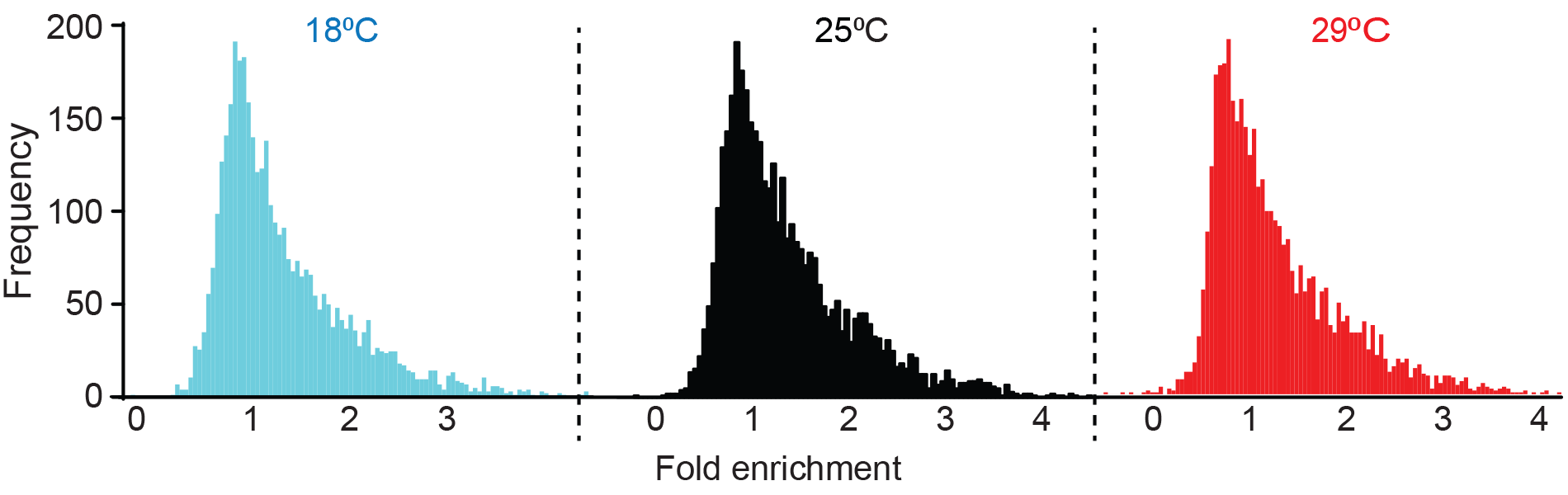
Density plot representing 4000 genes, selected as ‘AGO1-enriched/associated’. The distributions represent the frequency of each fold enrichment for a particular temperature.

**Figure S8.**
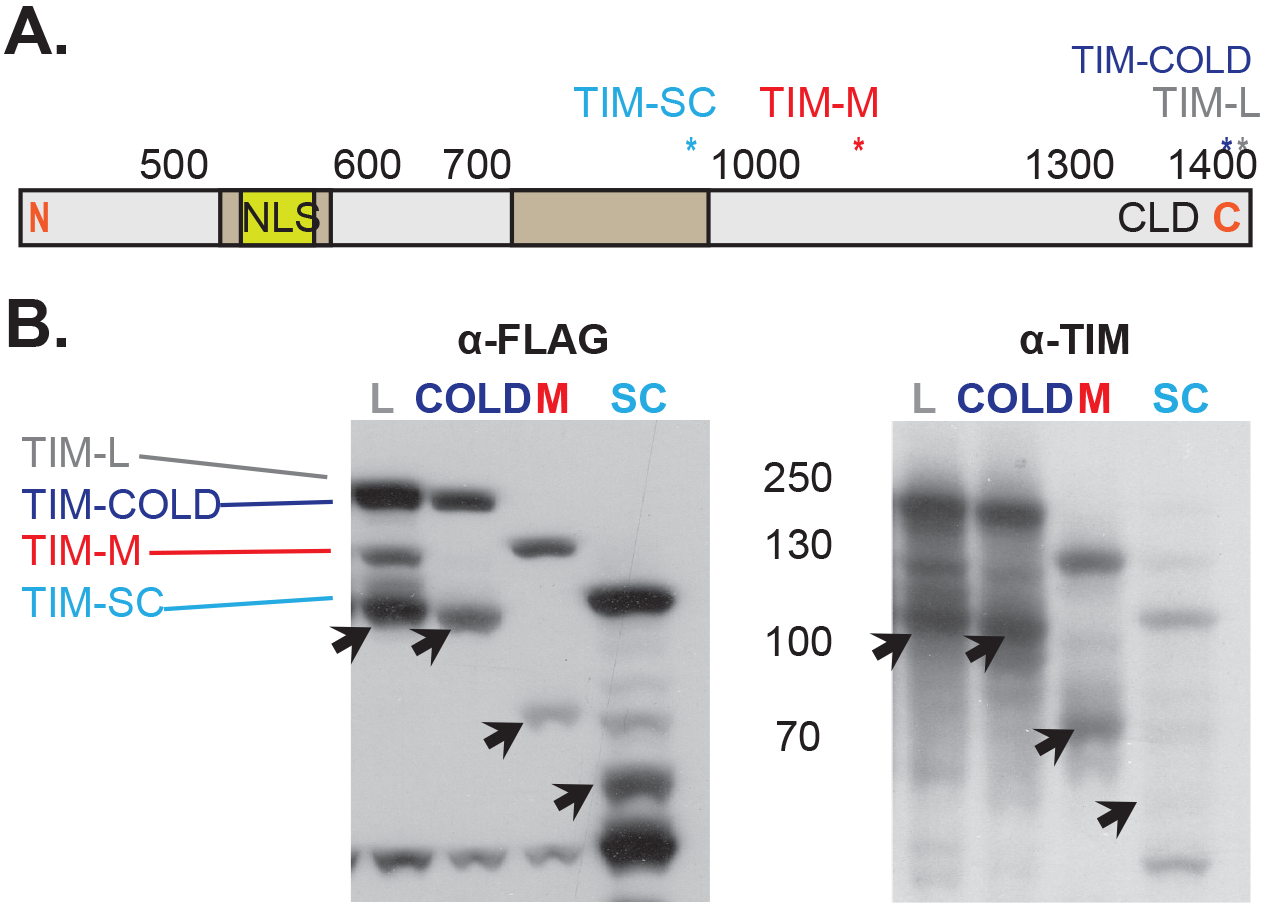
Overexpression of different *tim* isoforms with FLAG-tagged C-terminus. **A**. Schematic representation of TIM protein. The starts mark the location of the different FLAG-tags for each isoform. NLS: Nuclear Localization Signal. CLD: Cytoplasmic Localization Domain. **B.** Western blot membranes showing anti-FLAG and anti-TIM staining of the over-expression of the four 3’ FLAG-tagged UAS-*tim* isoforms in S2 cells. Arrows mark a degradation product ~ 60kDa smaller than the original TIM isoform.

**Figure S9.**
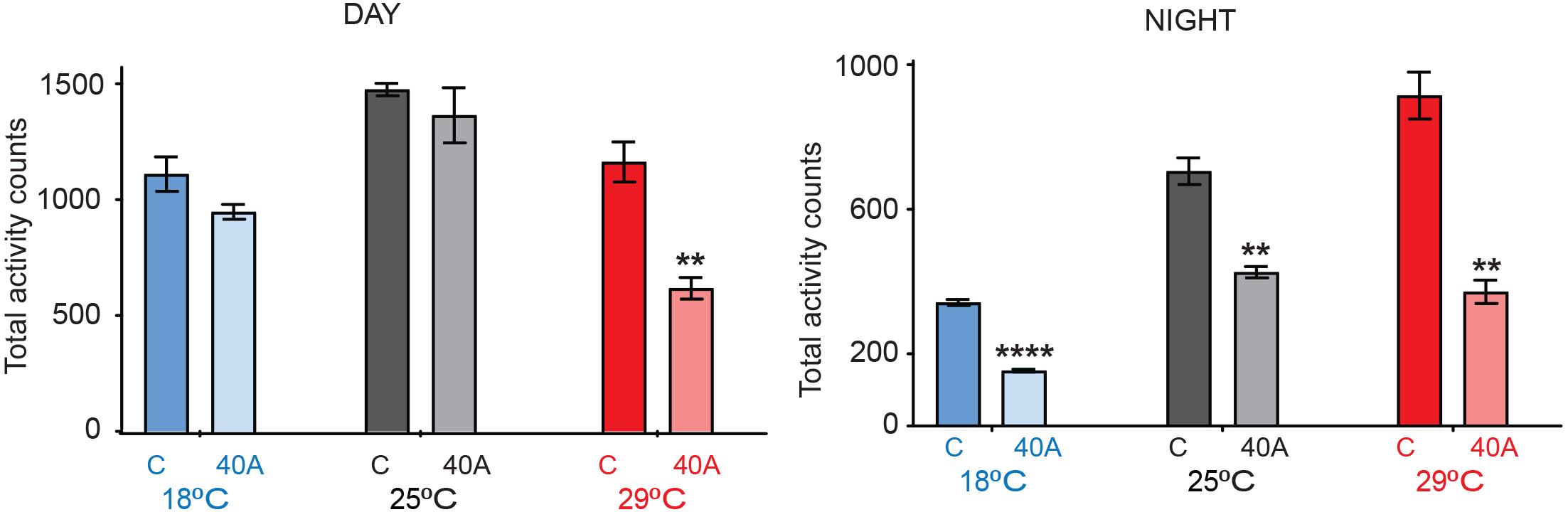
Total activity counts in control and 40A mutant flies at 18°C (blue), 25°C (grey) and 29°C(red) in LD12:12 during the day (left) and night (right). Stars represent significant differences between the control and mutants from the same temperature. Multiple t-test analysis using the Holm-Sidak method. **, pval<0.01 ***, pval<0.001. N=23−32.

**Figure S10.**
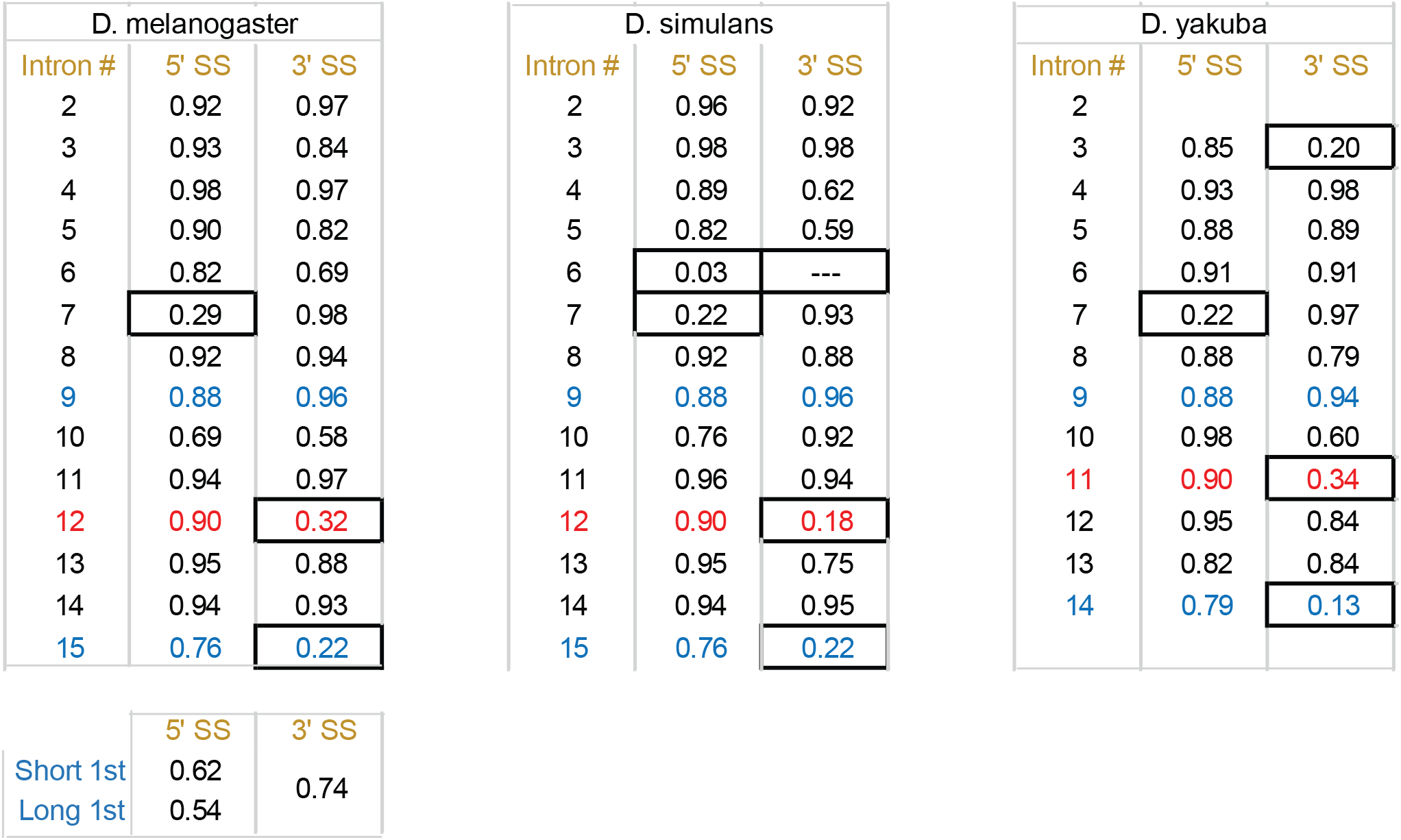
Summary table of the 3’ and 5’ splicing strengths of each *tim* intron for *Drosophila melanogaster, Drosophila simulans* and *Drosophila virilis*.

**Table S1. Circadian oscillation analysis.** The circadian analysis was performed using the MetaCycle algorithm (see main text for details). In the table the p-value, phase and amplitude are reported for each gene along with its normalized expression at each circadian timepoint.

**Table S2. Gene ontology analysis for the genes with circadian oscillation at each temperature.** Genes are ordered by ranking in Fisher exact test. Three different ontologies were analyzed: Biological process. Molecular function. Cellular component. Top 100 genes are reported.

**Table S3. Predicted miRNA binding sites (using TargetScan) for each *tim* RNA isoform 3’ UTR.** Putative miRNA binding sites are reported given the chromosome coordinates. Conservation in different insect species is also reported.

**Table S4. Differential miRNA expression at each temperature.** Log2(fold change) and p-adjusted value from DeSeq2 are reported for each miRNA.

